# SQMtools: automated processing and visual analysis of ’omics data with R and anvi’o

**DOI:** 10.1101/2020.04.23.057133

**Authors:** Fernando Puente-Sánchez, Natalia García-García, Javier Tamames

**Affiliations:** Systems Biology Department. Centro Nacional de Biotecnología (CNB-CSIC). C/ Darwin nº 3, Campus de Cantoblanco, 28049 Madrid, Spain

## Abstract

**Background:** The dramatic decrease in sequencing costs over the last decade has boosted the adoption of high-throughput sequencing applications as a standard tool for the analysis of environmental microbial communities. Nowadays even small research groups can easily obtain raw sequencing data. After that, however, non-specialists are faced with the double challenge of choosing among an ever-increasing array of analysis methodologies, and navigating the vast amounts of results returned by these approaches.

**Results:** Here we present a workflow that relies on the SqueezeMeta software for the automated processing of raw reads into annotated contigs and reconstructed genomes (bins). A set of custom scripts seamlessly integrates the output into the anvi’o analysis platform, allowing filtering and visual exploration of the results. Furthermore, we provide a software package with utility functions to expose the SqueezeMeta results to the R analysis environment.

**Conclusions:** Altogether, our workflow allows non-expert users to go from raw sequencing reads to custom plots with only a few powerful, flexible and well-documented commands.

## Background

The advent of high-throughput sequencing technologies in 2008 made it possible to directly sequence the different microbial genomes present in a given sample (metagenomics) as well as measuring the expression profiles of those genomes (metatranscriptomics) in a culture-independent manner. This supposed a revolution in the field of microbial ecology, as it allowed to profile the taxonomic composition and functional potential of microbial communities (Eisen, 2007) and eventually to recover full genomes from environmental bacteria that in some cases could not be studied by other means (Pedrós-Alió *et al.*, 2019).

Current sequencing costs (as of the end of 2019) are around 0.01$ per Megabase of DNA (Wetterstrand, 2019), which have led to the popularization of metagenomics as a powerful and affordable tool to analyze microbial communities. The democratization in the use of meta’omics is however hindered by the inherent complexity of high-throughput sequencing results, which need to be processed and analyzed by specialized personnel. Life science graduates often lack training in basic data science skills (Attwood *et al.*, 2017), and while large research groups might be able to hire one or more dedicated bioinformaticians, this is often not the case for smaller groups. There is thus a need for tools that allow life science researchers to reliably analyze meta’omics data with minimal training. Such tools should provide interfaces that relieve users from the burden of processing large amounts of data, while still giving them enough freedom to pose and answer biologically meaningful questions.

A standard metagenomics workflow comprises assembling shotgun reads into contigs, predicting genes on such contigs, taxonomically and functionally classifying contigs and genes, and estimating contig and gene abundances by remapping the shotgun reads against the assembly. Optionally, contigs can be combined into metagenome-assembled genomes (MAGs) or bins. This workflow is rather straightforward and thus amenable to be combined into a single pipeline that runs on sensible parameters with minimal user intervention. We recently implemented this concept in the SqueezeMeta pipeline for the automated analysis of metagenomes and metatranscriptomes (Tamames & Puente-Sánchez, 2019).

Once a metagenome has been processed, users face a second challenge, which is simply to navigate the vast amount of results. A standard-sized metagenome can contain from hundreds of thousands to millions of contigs, each coming from different organisms and containing different functions. Thus, even if results are already presented in a human-readable form (e.g. tables), their sheer size prevents non-experts from extracting the required information.

In this work, we seek to eliminate this problem by proposing an integrated workflow that takes care of all the steps of data processing and formatting, allowing users to easily propose and answer scientifically meaningful questions using their metagenomics/metatranscriptomics data. To achieve this, we here present a set of tools that seamlessly integrate the output of the SqueezeMeta pipeline with the anvi’o interactive visualization platform (Eren *et al.*, 2015) and the R environment for statistical analysis (R Core Team, 2013).

Anvi’o enables the interactive exploration of (meta)genomic data, and excels at bin refinement, but will struggle when trying to display a large number (i.e. millions) of contigs at the same time. For that, we provide a custom search engine that prefilters the data before launching the anvi’o interactive session. This engine allows users to input complex queries for selecting the contigs to be displayed based on their taxonomy, functional annotation and abundance across the different samples.

Additionally, we here describe the SQMtools R package, which automatically loads the different results generated by a SqueezeMeta run (e.g. contigs and gene sequences and annotations, aggregated functional and taxonomic profiles, binning results…) and exposes them as a single R object that can be then explored with a set of nine simple yet powerful functions. For example, users with no prior knowledge of the R programming language can easily generate a heatmap that displays the most abundant functions present in a taxa of interest across a set of samples, a barplot with the distribution of a given function across taxa in their metagenomes, or export a subset of their results into a spreadsheet. Furthermore, data loaded with the SQMtools package can be readily used with other popular analysis packages in microbial ecology, such as the DESeq2 package for differential abundance analysis (Love *et al.*, 2014), the vegan package for multivariate analysis (Oksanen *et al.*, 2007), or the caret package for machine learning (Kuhn, 2008).

## Results

To illustrate the utility and features of SQMtools, we used SqueezeMeta to analyze a set of 16 human gut metagenomes from two different populations with different lifestyles: urban-living Italian adult and hunter-gatherers from the Hadza people (**Supplementary Table S1**; Rampelli *et al.*, 2015). We were interested in identifying taxonomic and functional differences between the microbiomes of both populations. To that end, we loaded the data into R using the SQMtools package, and analysed it using the workflow illustrated in Figure 1 and detailed in the **Supplementary Material** (which includes all the code necessary to generate the figures in this article, and other examples showing additional features). We also show how to expose the data to other R packages such as vegan in order to perform more advanced analyses. Finally, we use anvi’o to visually explore subsets of the metagenomes, as well as the bins obtained by the SqueezeMeta pipeline.

**Figure 1.**
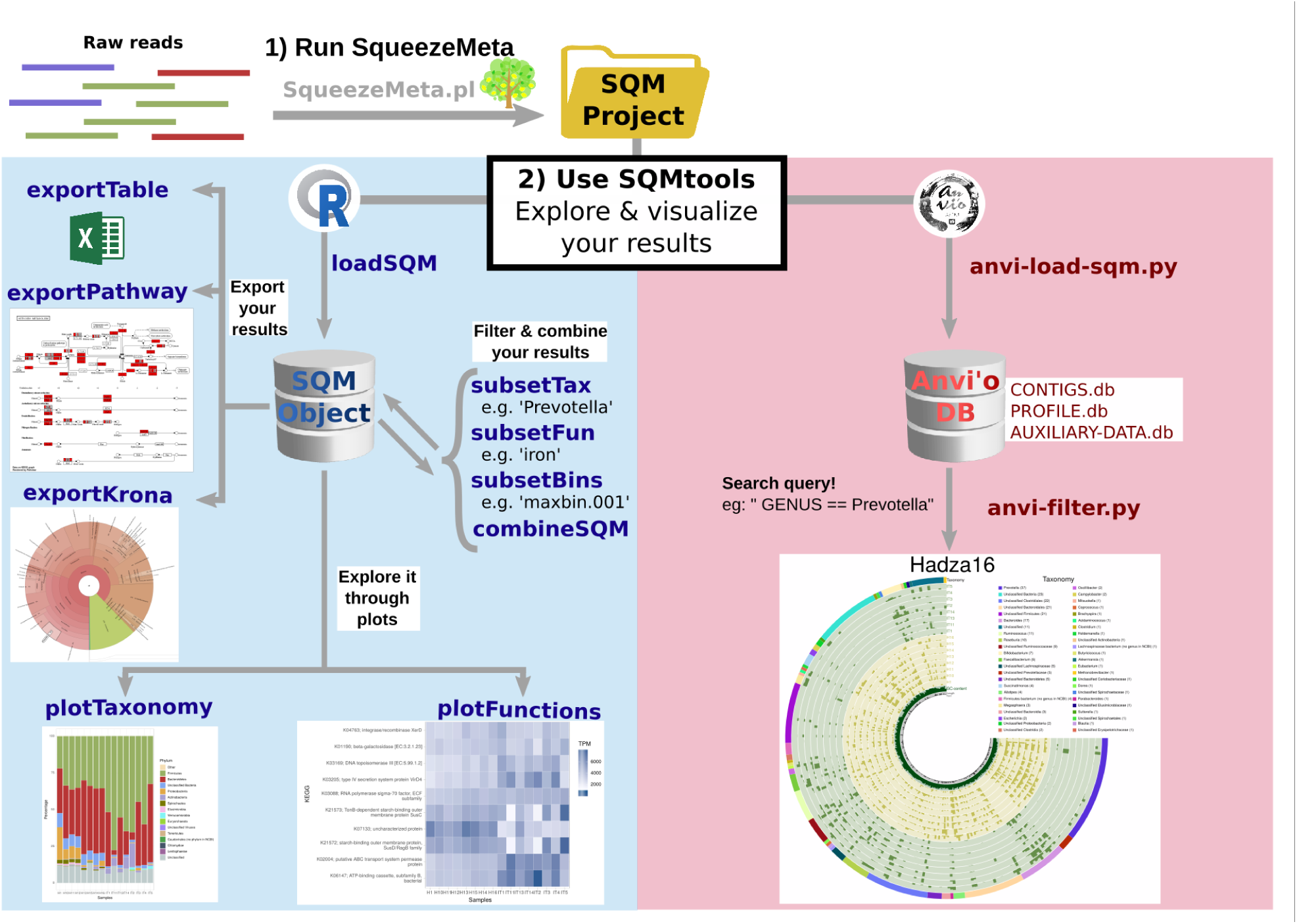
Main workflow of the SQMtools pipeline. Raw reads are processed automatically by SqueezeMeta, which integrates assembly, ORF prediction, annotation and binning. The results can then be easily filtered and explored either with R (by creating a SQM object) or anvi’o (by creating an anvi’o database) in order to produce custom plots with the information of interest. Arrows correspond to function calls (e.g. going from raw reads to a krona chart would involve three function calls). The workflow is exemplified in detail in the **Supplementary Information**.

Briefly the SQMtools contains the following types of functions:

- Load functions: load all the data generated by SqueezeMeta (sequences, annotations, bins, aggregated functional and taxonomic profiles) into a single R object (SQM object), whose structure is described in Supplementary Table S2, and which will be used by the rest of the main functions in the SQMtools package.
- Subset functions: generate a new SQM object containing a subset of functions, taxa, or bins of the parent SQM object.
- Combine functions: generate a new SQM object aggregating the data from two or more SQM objects.
- Plot functions: make different R plots from the data (taxonomy, functions…) contained in a SQM object.
- Export functions: create files (krona charts, KEGG pathway maps, tables…) from the data contained in a SQM object.

A general overview of the genus-level composition of the samples is presented in Figure 1a. Samples from Hadza individuals contain higher amounts of *Prevotella* (red), while samples from Italians contain higher amounts of *Bacteroides* (light blue). Non-metric dimensional scaling (NMDS) coupled to Permutational Multivariate Analysis of Variance shows that Hadza and Italian gut metagenomes can be clearly differentiated (*p* < 0.005) in terms of their taxonomic (Figure 2b) and functional (Figure 2c) compositions.

**Figure 2.**
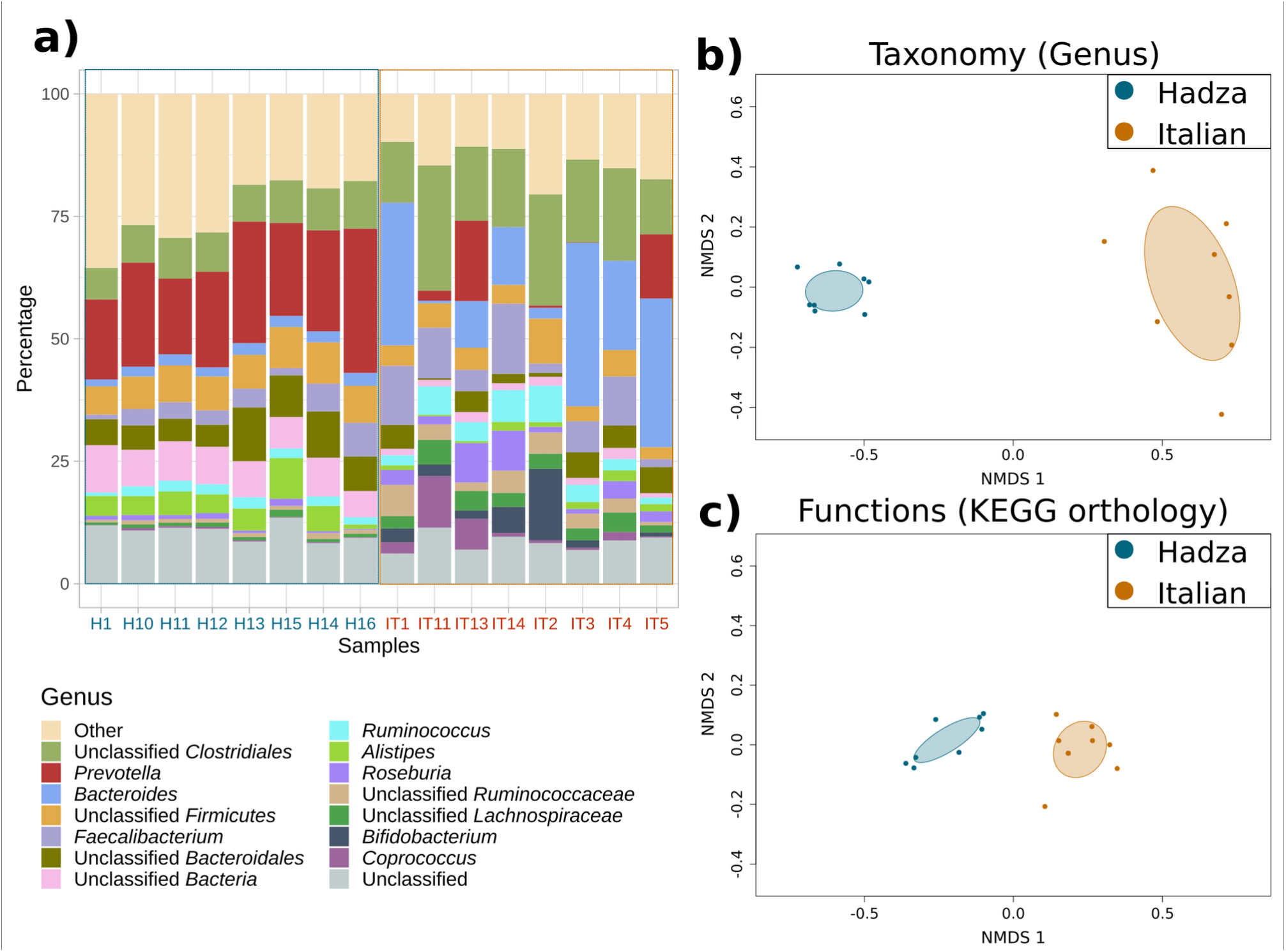
General overview of the example dataset. **a)** Genus-level taxonomic composition of the Hadza (blue) and Italian (orange) samples, as depicted by the SQMtools package. **b,c)** Nonmetric dimensional scaling (NMDS) showing the clustering of Hadza vs Italian samples based on their **b)** genus-level taxonomy and **c)** KEGG orthology functional profiling. Differences between clusters were significant (Permutational Multivariate Analysis of Variance *p* < 0.005).

We then show how SQMtools can be used to explore an individual metabolic pathway, in this case the biosynthesis of aromatic amino acids. Figure 3a shows the KEGG pathway map, with colours indicating whether a reaction was more present in the Hadza (blue) or Italian (orange) samples. As reported by Rampelli *et al*., (2015), dietary differences result in the Hadza microbiome being enriched in functions related to the biosynthesis of Phenylalanine and Tyrosine (Figure 3a, **box**), in particular the transaminase *tyrB* / K00832 / EC:2.6.1.57 catalysing the biosynthesis of the common intermediary L-arogenate. Figure 3b shows an anvi’o visualization obtained with the *anvi-filter.py* script, containing the taxonomy and distribution across samples of the contigs containing genes related to the biosynthesis of aromatic amino acids. The pathway is present in *Prevotella* for the Hadza samples (blue colored rings) and in *Bacteroides* for the Italian samples (orange colored rings), consistently with the taxonomic distribution presented in Figure 1a. Note that, when executed by the user, this visualization would appear in a browser window, and could be explored interactively. Finally, the abundance of individual KEGG orthology groups in each sample is presented as a heatmap in Figure 3c.

**Figure 3.**
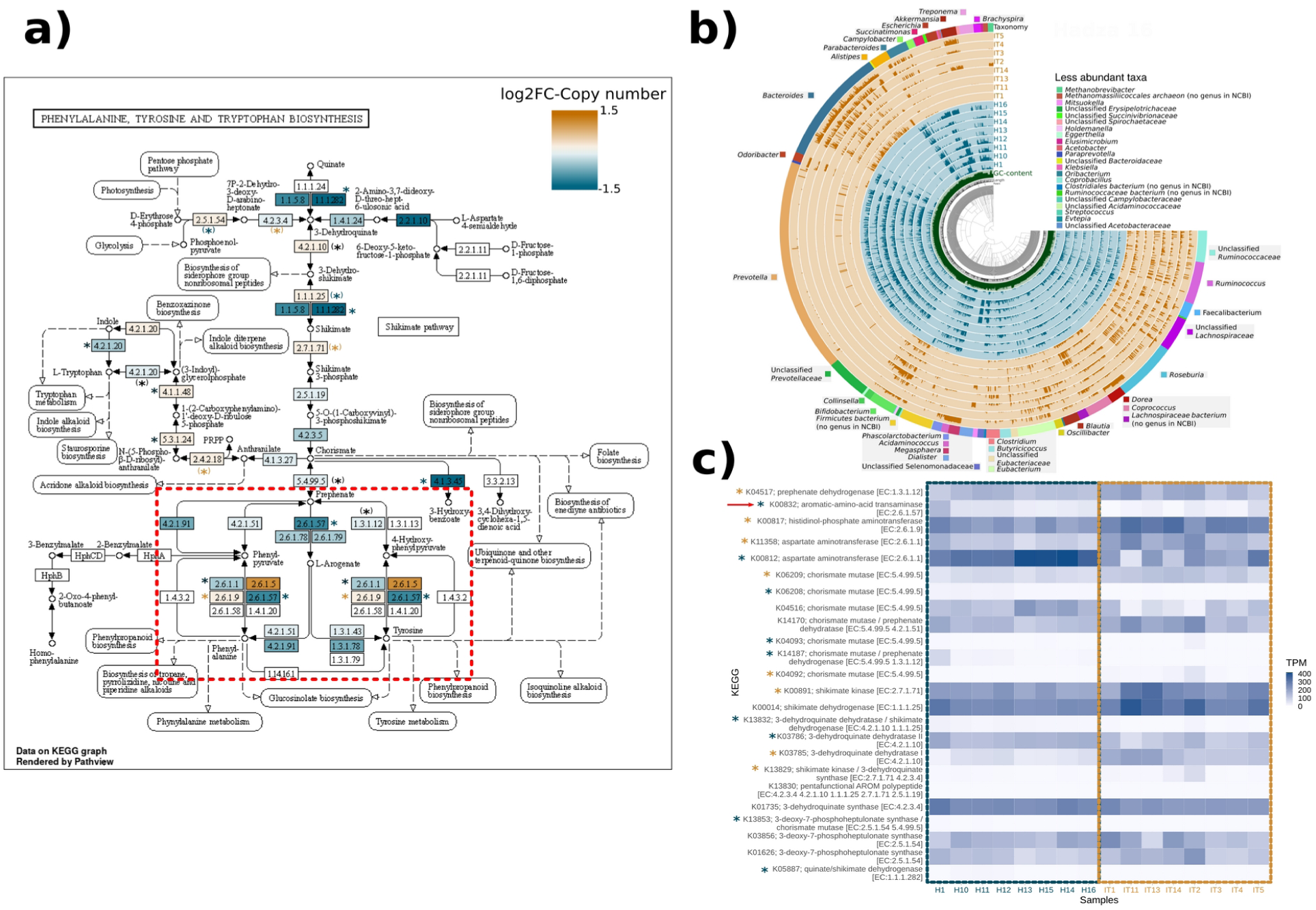
Detailed overview of the biosynthesis of aromatic amino acids. **a)** KEGG pathway map for Phenylalanine, Tyrosine and Tryptophan biosynthesis. Reactions are coloured based on the log2 fold-change of their average copy numbers in Italian (orange) vs Hadza (blue) samples. Asterisks indicated reactions performed by KEGG orthologs that were significantly enriched in Hadza (blue) or Italian (orange) samples (DESeq2 *p adj*. < 0.05). Black asterisks indicate reactions in which a different KEGG ortholog was enriched in Hadza and Italian samples (note that the same reaction can be performed by more than one KEGG ortholog). Asterisks within parentheses indicate reactions in which some, but not all KEGG orthologs were significantly enriched in a group. **b)** anvi’o plot showing the taxonomy and distribution across samples of the contigs containing genes related to the biosynthesis of aromatic amino acids. **c)** Heatmap depicting the TPM across samples of selected KEGG orthologs related to the biosynthesis of aromatic amino acids. Asterisks indicate significant enrichment in Hadza or Italian samples as described above.

Finally, Figure 4 shows the weight (in percentage of total reads) and taxonomic distribution of six broad functional categories across the different samples. Some taxa, like *Prevotella* (red) and *Bacteroides* (light blue), seem capable of metabolizing a large array of carbohydrates, while others, like *Bifidobacterium (*dark blue), seem for the most part limited to a few substrates.

**Figure 4.**
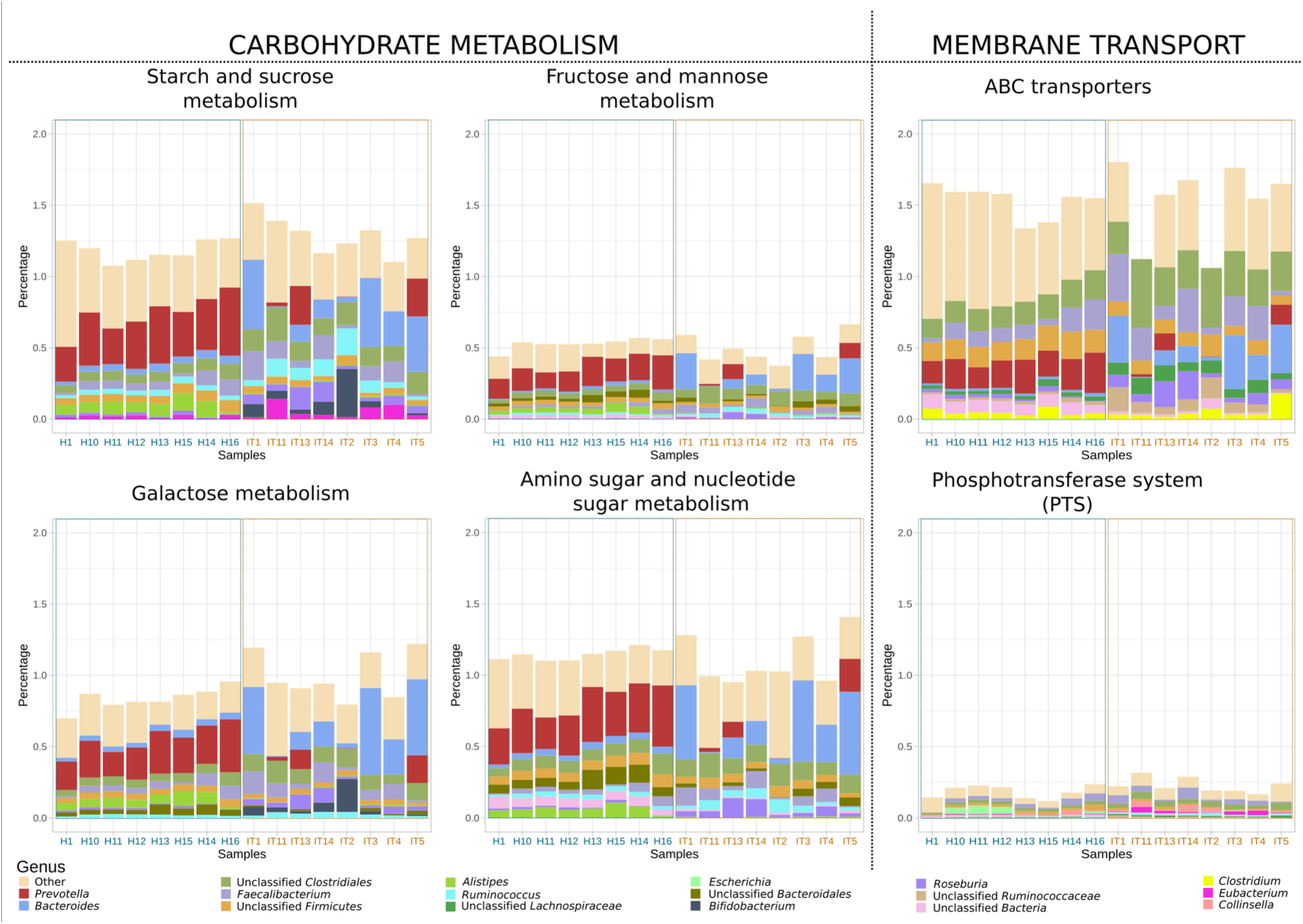
Taxonomic distribution of selected functions. Barplots showing the weight (in percentage of total reads) and taxonomic distribution of six broad functional categories across the different samples.

Overall, our results agree well with those presented by Rampelli *et al*., (2015). Save for minor post-processing additions (e.g. adding titles, coloured axes labels, etc) all the plots presented in Figures 2–4 can be generated by novice users with only a few commands, which illustrates the potential of this approach.

## Discussion

There are several software packages for the analysis and visualization of metagenomic data, with Galaxy (Goecks *et al.*, 2010), MEGAN (Huson *et al*., 2016) and STAMP (Parks *et al.*, 2014) being among the most popular. Our workflow differentiates from them in that 1) it covers all the steps of the analysis, from raw reads to custom figures and statistics and 2) it exposes the results to the R analysis environment, where they can be explored with a set of convenience functions included in the SQMtools package, but also directly analysed with other tools from the R ecosystem.

SQMtools provides different ways to look at the abundance of orfs, contigs, bins, taxa and functions in the metagenome (Table 1). In addition to read and base counts, which have a straightforward interpretation, we also provide TPM and copy numbers in order to take into account both sequencing depth and feature length. The TPM (transcripts per million) metric was introduced by Wagner *et al.*, (2012) as an improved way to account for gene length and sequencing depth in transcriptomic experiments: we find it equally useful in metagenomics. The TPM of a feature (be it a transcript, a gene or a functional category) is the number of times that we would find that feature when randomly sampling 1 million features, given the abundances of the different features in our sample. As an alternative to TPM, we also provide copy numbers, calculated as the ratio between the coverage of function of interest and the coverage of the RecA/RadA recombinase universal single-copy gene. By default, the *subsetTax* and *subsetBins* functions will rescale TPM and copy numbers so that they relate to the taxa present in the subset, rather than to the whole metagenome. Details on the interpretation of those values in the different subsets are shown in Table 1.

**Table 1.**
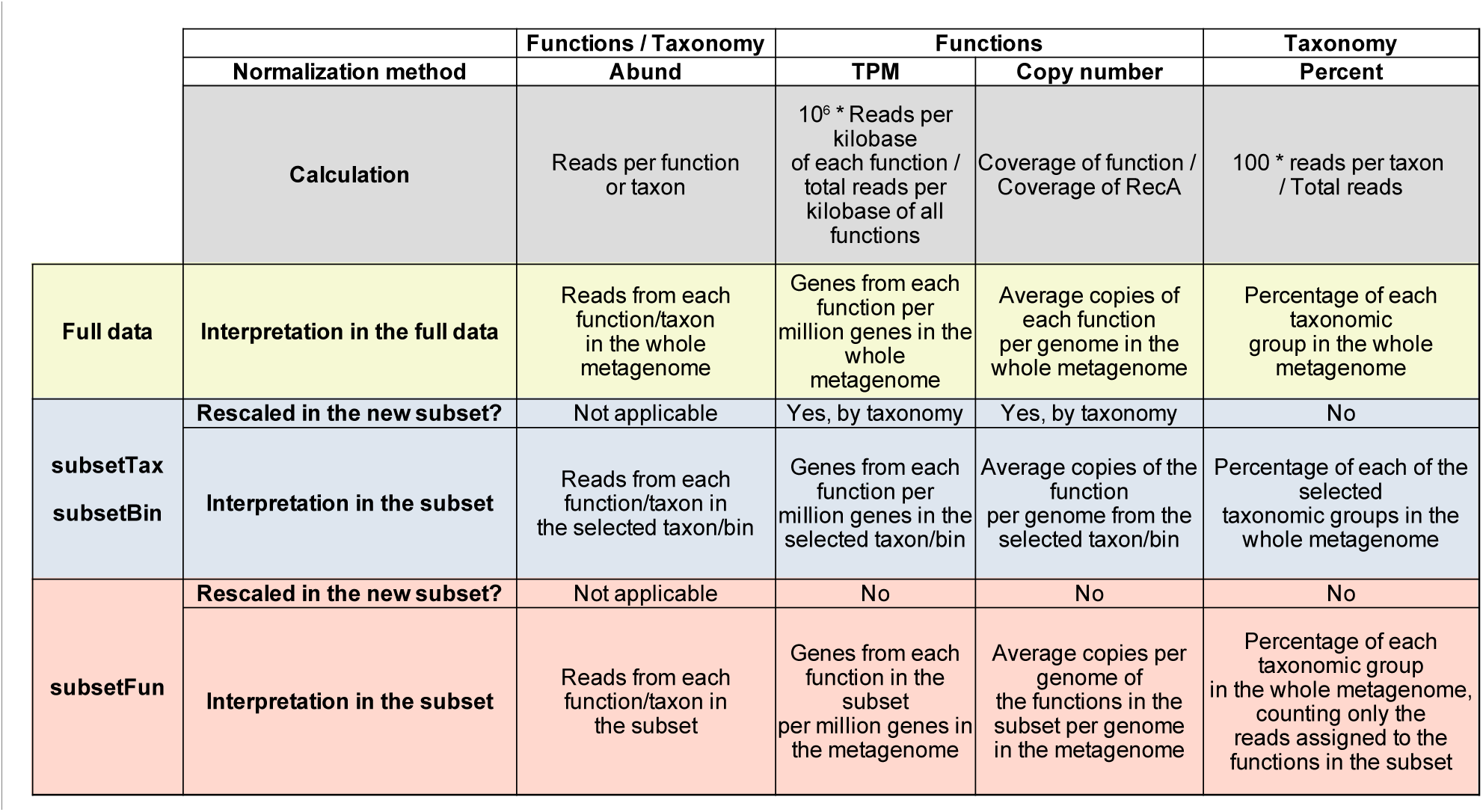
Abundance metrics included in SQMtools and their interpretation in the full metagenome and in taxonomic and functional subsets.

Note that TPMs are based on proportions calculated over an arbitrary count total (in this case, the number of reads sequenced per sample) and as such are compositional (see Gloor *et al*., 2017 for a detailed discussion on the problem and its implications). For visualization/exploration purposes, we often find it useful to compare TPMs from different samples (e.g. in Figure 3c) but in general statistical tests should be performed using raw reads and composition-aware methods (Gloor *et al*., 2017). Copy numbers, on the other hand, are ratios and should be less affected by compositionality issues (Morton *et al*. 2019). We nonetheless still recommend caution when performing statistical tests. The debate over the best way to analyze microbiome data is still ongoing (see e.g. two alternative avenues in Quinn *et al*., 2019 and Cruz *et al.*, 2020) and older methods such as DESeq2 might still outperform composition-aware methods for some applications (see **Methods**). In SQMtools, we provide common normalizations that can facilitate data exploration as well as the raw abundance data required to perform more advanced analyses, but we do not promote any particular statistical method as we expect the field to continue evolving.

## Conclusions

While many tools for metagenomics are available (Breitwieser *et al.*, 2019), to our knowledge we are the first to provide a robust, comprehensive and well-curated software suite covering all the steps of the analysis. This ensures full compatibility between the different included tools, and facilitates installation (a single instruction if using the conda package manager). Integration with R and anvi’o is effortless, relieving the users from the burden of parsing the complex output files that are common to metagenomic studies. Overall, our workflow allows non-expert users to go from raw sequencing reads to custom plots with only a few powerful, flexible and well-documented commands, while also facilitating the incorporation of more advanced statistical methods to their analyses.

## Methods

### Software implementation

The SqueezeMeta to anvi’o interface is implemented in python3, and consists of two scripts. Firstly, the *anvi-load-sqm.py* script will parse a whole SqueezeMeta project (annotated orfs, contigs and bins) into anvi’o (Eren *et al.*, 2015), generating databases that can be directly used with the *anvi-interactive* or *anvi-refine* tools from the anvi’o suite. Secondly, the *anvi-filter-sqm.py* integrates a search engine for prefiltering the data before launching the anvi’o interactive session. The SQM to R interface is implemented in the SQMtools R package, which contains several utility functions. The main components of the package are described in the next four sections. All analyses were performed using SqueezeMeta v1.1.1 and SQMtools v0.4.5.

### The SQM object

The SQM object is a custom R object (implemented as a S3 class) which contain all the relevant information from a metagenomic experiment (orf, contig and bin annotations, aggregated taxonomic and functional profiles, etc). The formal structure of the SQM object is defined in **Supplementary Table S2**. The *loadSQM* function from the SQMtools package will create a SQM folder by reading and parsing the results directory generated by the SqueezeMeta pipeline (Tamames & Puente-Sánchez, 2019). The SQM object can then be explored by a set of utility functions included in the SQMtools package (Figure 1). Alternatively, expert users can directly access its content in order to perform custom analyses.

### Normalization of sequencing data

For individual orfs, contigs and bins, we provide the following abundance metrics: mapped read count, mapped base counts, and TPM. The TPM (transcripts per million) metric was introduced by Wagner *et al.*, (2012) and its calculation is performed in two steps:

1. Obtain the RPK (reads per kilobase) of each feature, by dividing the number of mapped reads to that feature in that sample by the feature length in kilobases.
2. Calculate the TPM of each feature by dividing its RPK by the sum of RPKs of all the features in that sample, and multiplying by a million.

For the sake of being consistent with previous works, we maintain the nomenclature “TPM”, even when use it to measure the abundances of features other than transcripts.

For functional categories, we provide the following abundance metrics: mapped read counts, mapped base counts, TPM and copy number. The TPM of a given functional category is calculated as described above. The reads mapping to genes from that category are aggregated and divided by the average length of the genes from that category in the assembly. Copy number is calculated by dividing the aggregated coverage of the genes from that functional category by the coverage of COG0468 to the RecA/RadA recombinase, which is a universal single copy gene.

### The subset methods

The subset methods can be applied to every SQM object in order to generate smaller SQM objects containing the taxa or functions of interest. The *subsetTax* and *subsetBins* methods will select the contigs assigned to the requested taxon and bin, respectively, and the ORFs contained in those contigs. By default, they will rescale TPMs and copy numbers so that they are calculated with respect to the selected taxon/bins, rather than to the whole metagenome (Table 1).

The *subsetFun* method will directly select all the ORFs containing the requested functions. By default, functions are searched for by string matching against all the functional annotations (classification against KEGG, COG, PFAM and any other database provided by the user when running SqueezeMeta). Users can also search within the KEGG and COG functional hierarchies (e.g. Phenylalanine, Tyrosine and Tryptophan biosynthesis). Finally, users can search for functions using regular expressions instead of string matching by adding the parameter *fixed=F*.

### Figure generation

Heatmaps and barplots are generated using ggplot2. We also use KronaTools (Ondov *et al.*, 2011) for generating Krona charts, and pathview (Luo & Brouwer, 2013) for generating annotated KEGG pathway maps. Anvi’o plots were generated with anvi’o version 6.1 (Eren *et al.*, 2015).

### Multivariate and differential abundance analyses

Multivariate analysis was performed using the *metaMDS* and *adonis* functions from *vegan* version 2.5.6 (Oksanen *et al.*, 2007). Differential abundance analysis was performed with DESeq2 version (Love *et al.*, 2014) using an adjusted p-value cutoff of 0.05. DESeq2 was chosen in this study as it was shown to have a good precision/recall balance in simulated datasets with 5 and 10, replicates per group (Quinn *et al.*, 2019), which is close to our 8 replicates per cohort.

### Software availability

SqueezeMeta and SQMtools are available under a GPL3 license at https://github.com/jtamames/SqueezeMeta.

## Declarations

### Ethics approval and consent to participate

Not applicable

### Consent for publication

Not applicable

### Availability of data and materials

The raw datasets analysed during the current study are available in the NCBI repository, under BioProject PRJNA278393 (SRA Study: SRP056480; https://www.ncbi.nlm.nih.gov/sra/?term=SRP056480; Rampelli *et al.*, 2015).

### Authors’ contributions

FP-S and JT devised the package. FP-S designed and wrote the core SQMtools functions, FP-S and NG-G wrote the code involved in figure generation. All authors contributed to the SQM to anvi’o interface. FP-S and NG-G wrote the manuscript. All authors discussed the manuscript and approved the final version.

## Acknowledgements

The authors thank Giuseppe D’Auria (FISABIO) for the initial contribution of the SQMtools – KronaTools interface.

