## Supplementary Material for "SQMtools: automated processing and visual analysis of ’omics data with R and anvi’o"

---

Supplementary material

Fernando Puente-Sánchez [1,*] , Natalia García-García [1] and Javier Tamames [1]

[1] Systems Biology Department. Centro Nacional de Biotecnología (CNB-CSIC).

C/ Darwin no 3, Campus de Cantoblanco, 28049 Madrid, Spain.

[*]To whom correspondence should be addressed.

### 1. Running SqueezeMeta

Firstly, the user has to run SqueezeMeta over a set of raw samples. These samples can be provided with a tab-separated file (See **Table S1**). In this example, we have used a set of samples publicly available in the BioProject PRJNA278393 (SRA Study: SRP056480) 'Metagenome sequencing of the Hadza hunter-gatherer gut microbiota' (https://www.ncbi.nlm.nih.gov/bioproject/278393)[1].

**Table S1:** Samples file required by SqueezeMeta [1]

| | | |
|---|---|---|
| H1 | SRR1927149_1.fastq.gz | pair1 |
| H1 | SRR1927149_2.fastq.gz | pair2 |
| H10 | SRR1929408_1.fastq.gz | pair1 |
| H10 | SRR1929408_2.fastq.gz | pair2 |
| H11 | SRR1929484_1.fastq.gz | pair1 |
| H11 | SRR1929484_2.fastq.gz | pair2 |
| H12 | SRR1929485_1.fastq.gz | pair1 |
| H12 | SRR1929485_2.fastq.gz | pair2 |
| H13 | SRR1929563_1.fastq.gz | pair1 |
| H13 | SRR1929563_2.fastq.gz | pair2 |
| H15 | SRR1929574_1.fastq.gz | pair1 |
| H15 | SRR1929574_2.fastq.gz | pair2 |
| H14 | SRR1930121_1.fastq.gz | pair1 |
| H14 | SRR1930121_2.fastq.gz | pair2 |
| H16 | SRR1930122_1.fastq.gz | pair1 |
| H16 | SRR1930122_2.fastq.gz | pair2 |
| IT1 | SRR1930247_1.fastq.gz | pair1 |
| IT1 | SRR1930247_2.fastq.gz | pair2 |
| IT11 | SRR1930248_1.fastq.gz | pair1 |
| IT11 | SRR1930248_2.fastq.gz | pair2 |
| IT13 | SRR1930251_1.fastq.gz | pair1 |
| IT13 | SRR1930251_2.fastq.gz | pair2 |
| IT14 | SRR1930250_1.fastq.gz | pair1 |
| IT14 | SRR1930250_2.fastq.gz | pair2 |
| IT2 | SRR1930253_1.fastq.gz | pair1 |
| IT2 | SRR1930253_2.fastq.gz | pair2 |
| IT3 | SRR1930255_1.fastq.gz | pair1 |
| IT3 | SRR1930255_2.fastq.gz | pair2 |
| IT4 | SRR1930777_1.fastq.gz | pair1 |
| IT4 | SRR1930777_2.fastq.gz | pair2 |
| IT5 | SRR1931170_1.fastq.gz | pair1 |
| IT5 | SRR1931170_2.fastq.gz | pair2 |

[1] Tab-separated file in which the first column shows sample names, the second contains the name of the fastq files: <SRA Accession Number> plus '_1' for forward reads and '_2' for reverse reads, and the third is the pair number of the reads.

```
SqueezeMeta.pl -m coassembly -p Hadza16 -s Hadza16.samples -f raw
```

where: '-m' is the assembly mode; '-p', the project name; '-s', the samples file; and '-f', the Fastq read files' directory (see the SqueezeMeta repository: https://github.com/jtamames/SqueezeMeta).
After running SqueezeMeta, we proposed two approaches based on R and anvi'o environments to explore the results.

[1]Rampelli, *et al.*. **(2015)**. Metagenome sequencing of the Hadza hunter-gatherer gut microbiota. *Current Biology, 25(13), 1682-1693).*

### 2. Exploring a SqueezeMeta project in R

#### 2.1. Loading a SqueezeMeta project in R

We propose the SQMtools R package to easily handle the SqueezeMeta output.

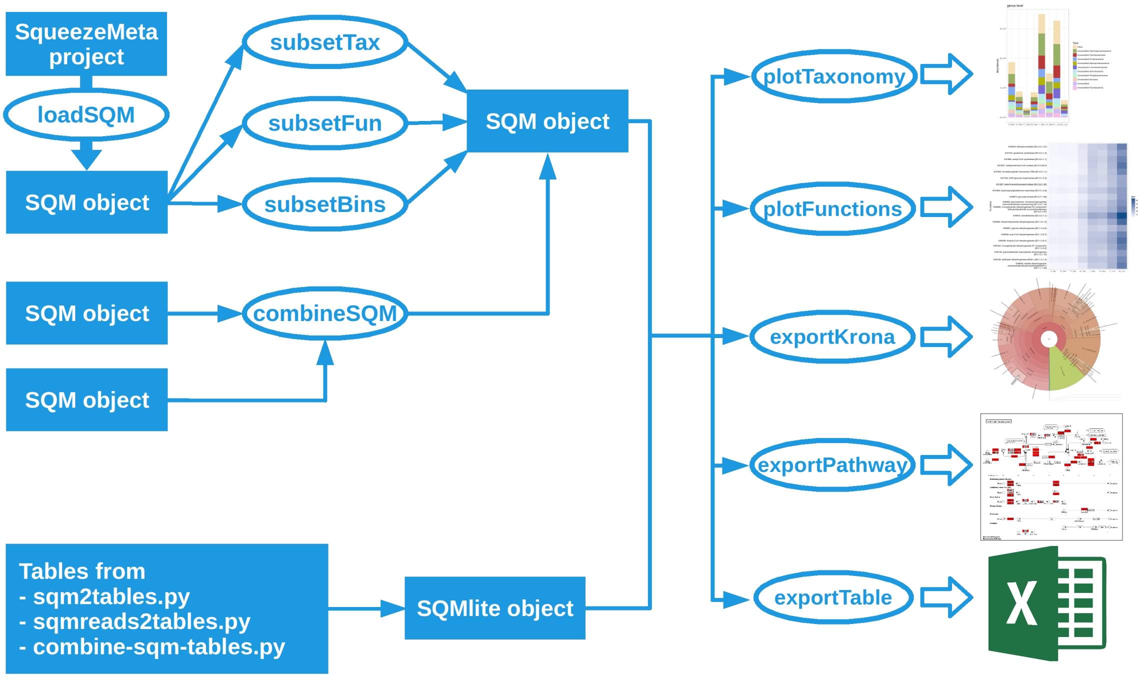

**Figure S1:** R workflow using *SQMtools*.

Firstly, the SqueezeMeta project must be loaded in R using the *loadSQM* function:

```
library('SQMtools')
Hadza = loadSQM('Hadza16')
```

This function creates a single SQM object that gathers all the data from SqueezeMeta.

**Table S2:** R SQM object structure.

| | lvl1 | lvl2 | lvl3 | type | rows/names | columns | data |
|---|---|---|---|---|---|---|---|
| SQM | $orfs | $table | | dataframe | orfs | misc. data | misc. data |
| | | $abund | | numeric matrix | orfs | samples | abundances (reads) |
| | | $bases | | numeric matrix | orfs | samples | abundances (bases) |
| | | $tpm | | numeric matrix | orfs | samples | tpm |
| | | $seqs | | character vector | orfs | (n/a) | sequences |
| | | $tax | | character matrix | orfs | tax. ranks | taxonomy |
| | $contigs | $table | | dataframe | contigs | misc. data | misc. data |
| | | $abund | | numeric matrix | contigs | samples | abundances (reads) |
| | | $tpm | | numeric matrix | contigs | samples | tpm |
| | | $seqs | | character vector | contigs | (n/a) | sequences |
| | | $tax | | character matrix | contigs | tax. ranks | taxonomies |
| | | $bins | | character matrix | contigs | bin. methods | bins |
| | $bins | $table | | dataframe | bins | misc. data | misc. data |
| | | $tpm | | numeric matrix | bins | samples | tpm |
| | | $tax | | character matrix | bins | tax. ranks | taxonomy |
| | $taxa | $superkingdom | $abund | numeric matrix | superkingdoms | samples | abundances (reads) |
| | | | $percent | numeric matrix | superkingdoms | samples | percentages |
| | | $phylum | $abund | numeric matrix | phyla | samples | abundances (reads) |
| | | | $percent | numeric matrix | phyla | samples | percentages |
| | | $class | $abund | numeric matrix | classes | samples | abundances (reads) |
| | | | $percent | numeric matrix | classes | samples | percentages |
| | | $order | $abund | numeric matrix | orders | samples | abundances (reads) |
| | | | $percent | numeric matrix | orders | samples | percentages |
| | | $family | $abund | numeric matrix | families | samples | abundances (reads) |
| | | | $percent | numeric matrix | families | samples | percentages |
| | | $genus | $abund | numeric matrix | genera | samples | abundances (reads) |
| | | | $percent | numeric matrix | genera | samples | percentages |
| | | $species | $abund | numeric matrix | species | samples | abundances (reads) |
| | | | $percent | numeric matrix | species | samples | percentages |
| | $functions | $KEGG | $abund | numeric matrix | KEGG ids | samples | abundances (reads) |
| | | | $bases | numeric matrix | KEGG ids | samples | abundances (bases) |
| | | | $tpm | numeric matrix | KEGG ids | samples | tpm |
| | | | $copy_number | numeric matrix | KEGG ids | samples | avg. copies |
| | | $COG | $abund | numeric matrix | COG ids | samples | abundances (reads) |
| | | | $bases | numeric matrix | COG ids | samples | abundances (bases) |
| | | | $tpm | numeric matrix | COG ids | samples | tpm |
| | | | $copy_number | numeric matrix | COG ids | samples | avg. copies |
| | | $PFAM | $abund | numeric matrix | PFAM ids | samples | abundances (reads) |
| | | | $bases | numeric matrix | PFAM ids | samples | abundances (bases) |
| | | | $tpm | numeric matrix | PFAM ids | samples | tpm |
| | | | $copy_number | numeric matrix | PFAM ids | samples | avg. copies |
| | $total_reads | | | numeric vector | samples | (n/a) | total reads |
| | $misc | $project_name | | character vector | (empty) | (n/a) | project name |
| | | $samples | | character vector | (empty) | (n/a) | samples |
| | | $tax_names_long | $superkingdom | character vector | short taxa names | (n/a) | long taxa names |
| | | | $phylum | character vector | short taxa names | (n/a) | long taxa names |
| | | | $class | character vector | short taxa names | (n/a) | long taxa names |
| | | | $order | character vector | short taxa names | (n/a) | long taxa names |
| | | | $family | character vector | short taxa names | (n/a) | long taxa names |
| | | | $genus | character vector | short taxa names | (n/a) | long taxa names |
| | | | $species | character vector | short taxa names | (n/a) | long taxa names |
| | | $tax_names_short | | character vector | long taxa names | (n/a) | short taxa names |
| | | $KEGG_names | | character vector | KEGG ids | (n/a) | KEGG names |
| | | $KEGG_paths | | character vector | KEGG ids | (n/a) | KEGG hierarchy |
| | | $COG_names | | character vector | COG ids | (n/a) | COG names |
| | | $COG_paths | | character vector | COG ids | (n/a) | COG hierarchy |
| | | $ext_annot_sources | | character vector | (empty) | (n/a) | External databases |

This object is a list, so data can be accessed using the syntax:

data_of_interest = <SQMobject>$<lvl1>$<lvl2>$<lvl3>.

```r
# Matrix genera vs. abundances
genus_tax = Hadza$taxa$genus$abund
# Show the first 5 rows and 5 columns
genus_tax[1:5, 1:5]
```

```
                                          H1 H10 H11 H12 H13
Unclassified Candidatus Aenigmarchaeota 46  13   5   1   0
Unclassified Candidatus Altiarchaeota    0   0   0   0   2
Unclassified Candidatus Bathyarchaeota  10 238  74  34  17
Unclassified Candidatus Korarchaeota     8   0   0   0   0
Unclassified Candidatus Lokiarchaeota    4  30   8   5  24
```

```
# Matrix COGs vs. abundances
COG_table = Hadza$functions$COG$abund
COG_table[1:5, 1:5]
```

```
           H1    H10   H11  H12  H13
COG0001 2560  9376 2613 1382 2306
COG0002 7475 14024 3281 2022 6014
COG0003 48    95    22    3    4
COG0004 1446  4754 1111  652 1670
COG0005 3724 10696 2625 1592 5021
```

```
# Raw table from SqueezeMeta: 13.<projectName>.orftable
ORF_table = Hadza$orfs$table
ORF_table[1:5, 1:5]
```

```
                  Contig ID  Molecule  Method   Length NT   Length AA
megahit_1_116-250 megahit_1      CDS   Prodigal       136         46
megahit_2_3-284   megahit_2      CDS   Prodigal       283         95
megahit_3_2-268   megahit_3      CDS   Prodigal       268         90
megahit_4_1-261   megahit_4      CDS   Prodigal       262         88
megahit_5_3-620   megahit_5      CDS   Prodigal       619        207
```

It is also possible to make a summary of the SqueezeMeta project:

```
smry_Hadza = summary(Hadza)
```

```
smry_Hadza$project_name
```

```
'Hadza16'
```

```
smry_Hadza$samples
```

```
'H1' 'H10' 'H11' 'H12' 'H13' 'H15' 'H14' 'H16' 'IT1' 'IT11' 'IT13' 'IT14' 'IT2' 'IT3'
'IT4' 'IT5'
```

```
smry_Hadza$reads_in_orfs
```

```
    H1       H10       H11       H12       H13       H15       H14       H16
    23847271 52692441 13718056  7833832 23807939 16412877 61547031 25641150
    IT1      IT11      IT13      IT14       IT2       IT3       IT4       IT5
    3191223   7090403   3415890   2472831 15259976 12563001  4362936 29855475
```

#### 2.2. Exploring and visualising the SqueezeMeta results

In this point, we show how to explore the newly loaded SqueezeMeta project through custom plots. They are a straightforward way to answer some of the most common ecological questions:

##### 2.2.1. Who is there? The *plotTaxonomy* function

This function selects the most abundant taxa across all samples in a SQM object and represents their abundances (counts/percents) in a barplot.

```
plotTaxonomy(Hadza, rank = 'phylum', count = 'percent')
```

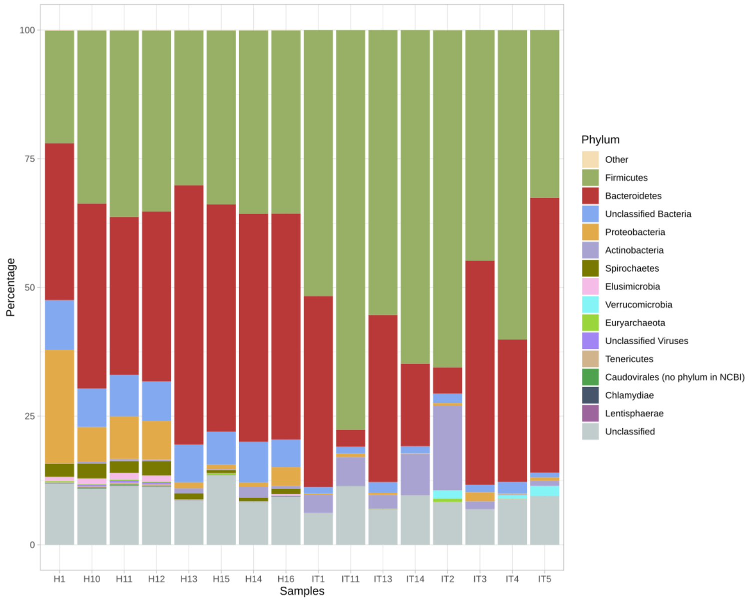

**Figure S2:** Taxonomy barplot at the phylum level. The abundance of each taxonomic group in the whole metagenome is shown in percentages.

```
plotTaxonomy(Hadza, rank = 'family', count = 'percent')
```

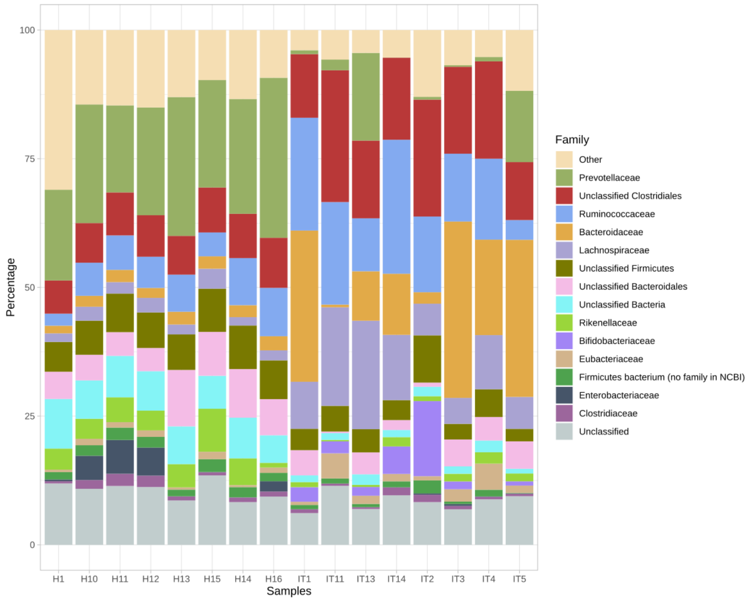

**Figure S3:** Taxonomy barplot at the family level.

```r
plotTaxonomy(Hadza, rank = 'genus', count = 'percent')
```

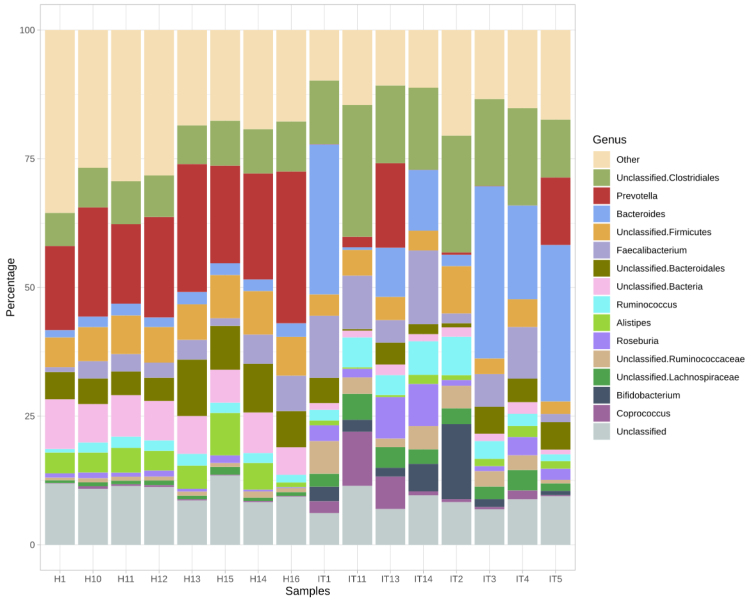

**Figure S4:** Taxonomy barplot at the genus level.

Alternatively, this function allows the user to customize the plots. Some of the options are: choosing colors, scaling percentages, picking up the most abundant or the user's favourite taxa, plotting or not the 'Unclassified' (features, contigs or genes, that have not been taxonomically classified) and the 'Other' category ('Other' is an artificial taxon that groups less abundant taxa).

```r
# Choosing taxa without re-scaling and without 'Other' taxon
plotTaxonomy(Hadza, rank = 'genus', count = 'percent', tax = c('Prevotella',
'Alistipes', 'Bacteroides', 'Bifidobacterium'), rescale = F, color = c('yellowgreen',
'darkolivegreen1', 'springgreen1', 'deepskyblue2'), others = F)
```

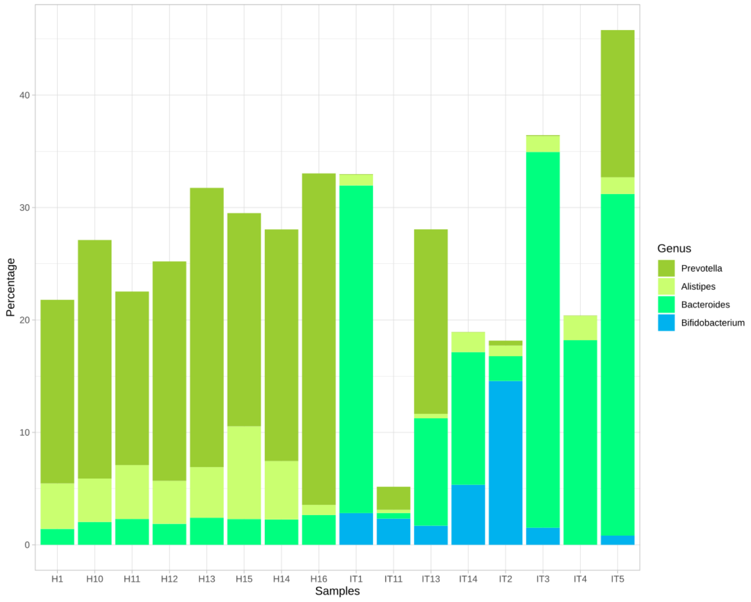

**Figure S5:** Taxonomy barplot of *Prevotella*, *Alistipes*, *Bacteroides* and *Bifidobacterium* genera without re-scaling percentages and without including the 'Other' taxon.

```
# Choosing taxa, re-scaling
plotTaxonomy(Hadza, rank = 'genus', count = 'percent', tax = c('Prevotella',
'Alistipes', 'Bacteroides', 'Bifidobacterium'), rescale = T, color = c('yellowgreen',
'darkolivegreen1', 'springgreen1', 'deepskyblue2'), others = T)
```

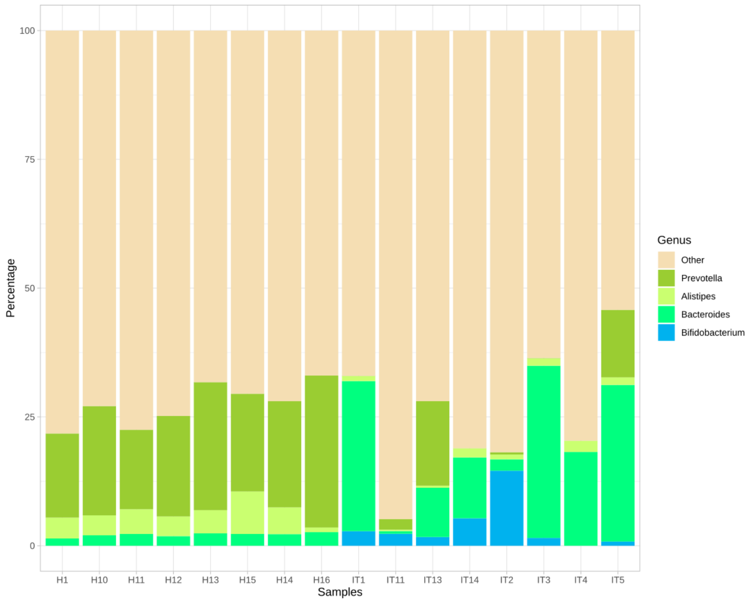

**Figure S6:** Taxonomy barplot of *Prevotella*, *Alistipes*, *Bacteroides* and *Bifidobacterium* genera re-scaling percentages and including the 'Other' taxon.

##### 2.2.2. What are they doing? The *plotFunctions* function

This function shows the most abundant functional categories (annotated with KEGGs, COGs and/or PFAMs) across all the samples in a SQM object and represents their abundances (counts, TPM or copy number) in a heatmap. Again, a custom set of functions can be represented.

```
plotFunctions(Hadza, fun_level = 'KEGG', count = 'tpm', N = 15)
```

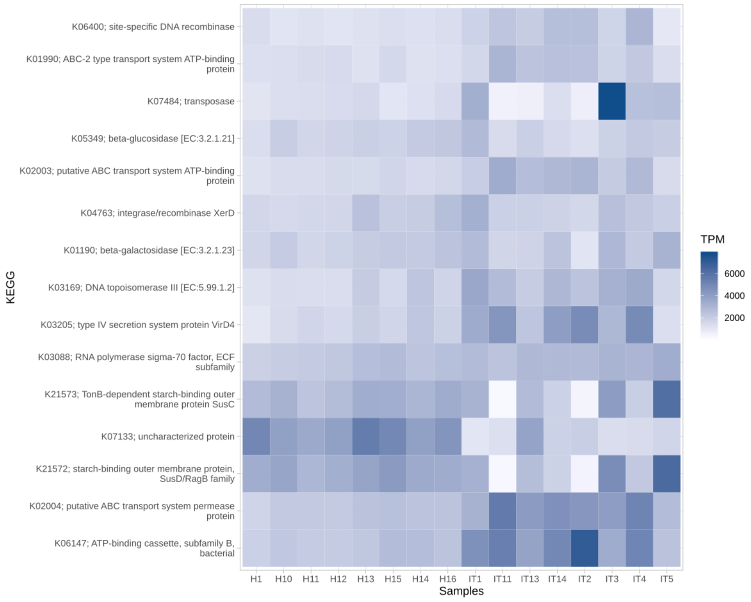

**Figure S7:** Most abundant KEGGs in the Hadza16 project. The abundance of each function (KEGG) is counted in TPM ('transcripts per million'), genes from each function per million genes in the whole metagenome.

```r
plotFunctions(Hadza, fun_level = 'KEGG', count = 'copy_number', N = 15)
```

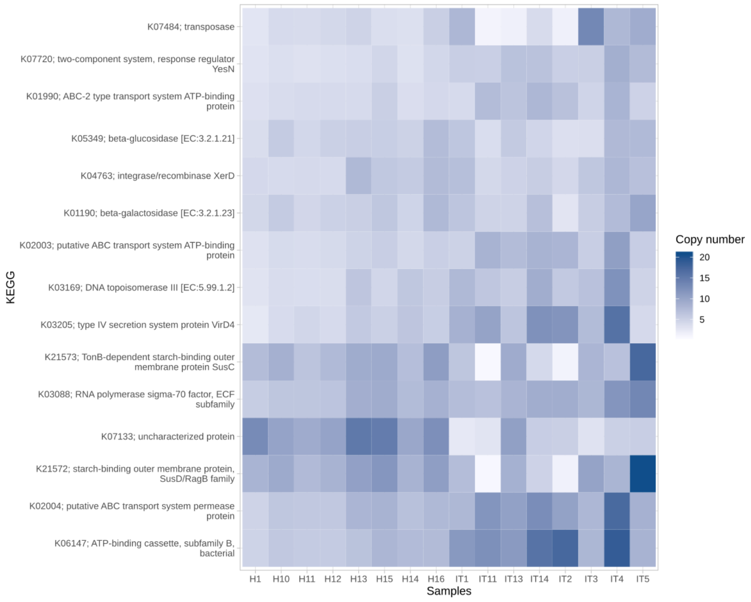

**Figure S8:** Most abundant KEGGs in the Hadza metagenome. The abundance of each function is counted in copy number (the average copies of each function per genome in the whole metagenome).

##### 2.2.3. Subsetting and combining results: The *subsetTax*, *subsetFun*, *subsetBins* and *combineSQM* functions

This package includes other functions designed to subset a SQM object based on specific taxa, functions, bins and, also, combine these reduced SQM objects. Since these new objects are also SQM objects, most of the functions of the package are still usable.

**Studying a specific set of functions.** E.g.: Phenylalanine, tyrosine and tryptophan biosynthesis.
If the users are interested in analyzing differences between Hadza e Italian gut microbiomes in the Phenylalanine, tyrosine and tryptophan biosynthesis, they can do a subset of all the functions related to it:

```r
aromatic_aa = subsetFun(Hadza, fun =
                'Phenylalanine, tyrosine and tryptophan biosynthesis')
# Plot taxonomy
plotTaxonomy(aromatic_aa, 'genus', 'percent', N = 10, rescale = F, others = T)
```

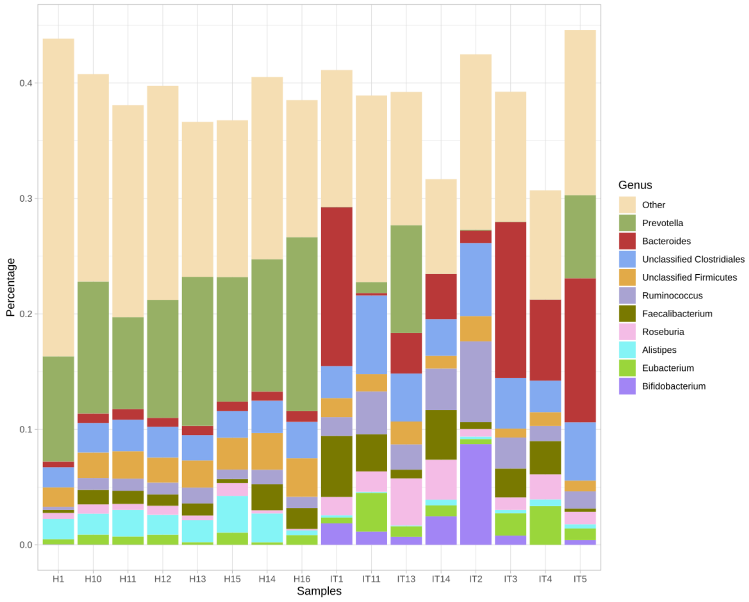

**Figure S9:** Taxonomy barplot at the genus level of the 'aromatic_aa' subset. The barplot shows the taxonomic distribution and percentage in the different samples of the ORFs containing functions from the 'Phenylalanine, tyrosine and tryptophan biosynthesis' KEGG pathway.

```
# Plot functions
plotFunctions(aromatic_aa, fun_level = 'KEGG', count = 'tpm')
```

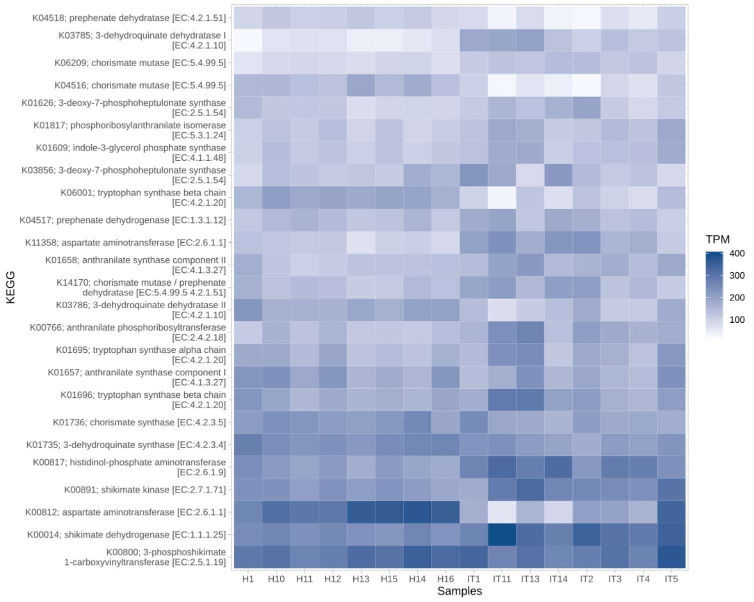

**Figure S10:** Most abundant KEGGs in the 'aromatic_aa' subset. The abundance of each function (KEGG) is counted in TPM ('transcripts per million'), genes from each function per million genes in the whole metagenome.

Specific KEGGs, COGs, PFAMs can be analyzed with more detail, for instance, extracting their sequence:

```
# Extract the ORF annotated with KEGG 'K00812'
K00812 = subsetFun(Hadza, 'K00812')
# Extract the sequence
sequence = K00812$orfs$seqs
sequence[1]
```

```
megahit_714_4687-5880
'MTKLSDRLNRLAPSATLAMSQKSGEMKAQGIDVINMSVGEPDFNTPDHIKEAAKKAIDENYSRYSPVPGYIELRKAIVE
KLKKENGLEYSTNEILVSNGAKQSVCNTVMALVNDGEEVIIPAPYWVSYPQMVKLAGGEPVIVEAGFEQNFKMTPEQLEAA
ITPKTRMLILCSPSNPTGSVYSEEELRGLAEVIKRHEGLYVLADEIYEHINYVGKHSSIAHIEGMRERTIIVNGVSKAYAMTGW
RIGFIAAPEWIVKGCNKLQGQYTSGPCSVSQKAAETAYTTSQECVETMRKAFERRRDLIVELAKDIPGLEVNKPEGAFYLFPKC
SSFFGKSYNGKKIENSTDLALFLLEVGHVATVGGDAFGDPYCFRMSYATSDDNIRKAMKRIKDTLALLA*'
```

**Exploring a specific taxon.** E.g. Contigs assigned to the '*Prevotella*' genus.

```
# Subset of Prevotella contigs
prevotella = subsetTax(Hadza, 'genus', tax = 'Prevotella')
# Plot taxonomy
plotTaxonomy(prevotella, 'genus',  'percent', N = 10, rescale = F, others = T)
```

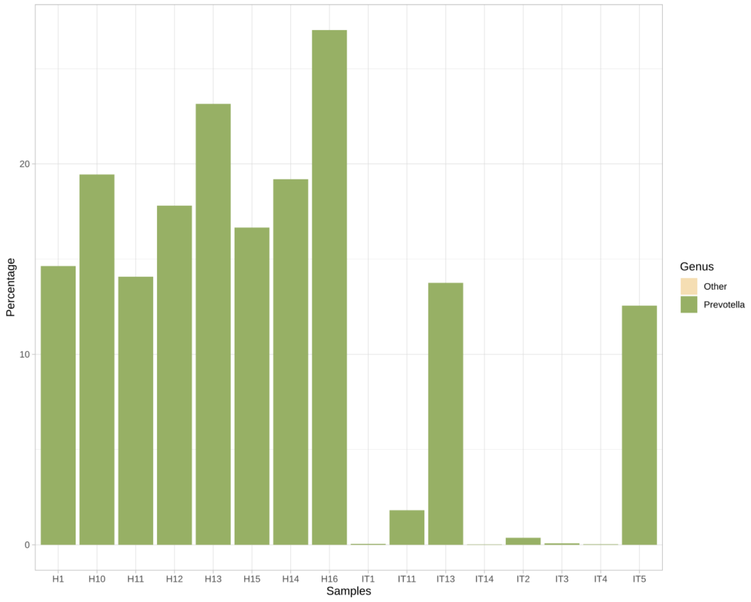

**Figure S11:** Taxonomy barplot at the genus level of the 'prevotella' subset. The abundance percentage of *Prevotella* is relative to the whole metagenome. Note that the abundance of *Prevotella* is very low in the samples IT1, IT11, IT14, IT2, IT3 and IT4.

```
# Plot functions
plotFunctions(prevotella, fun_level = 'KEGG', count = 'tpm')
```

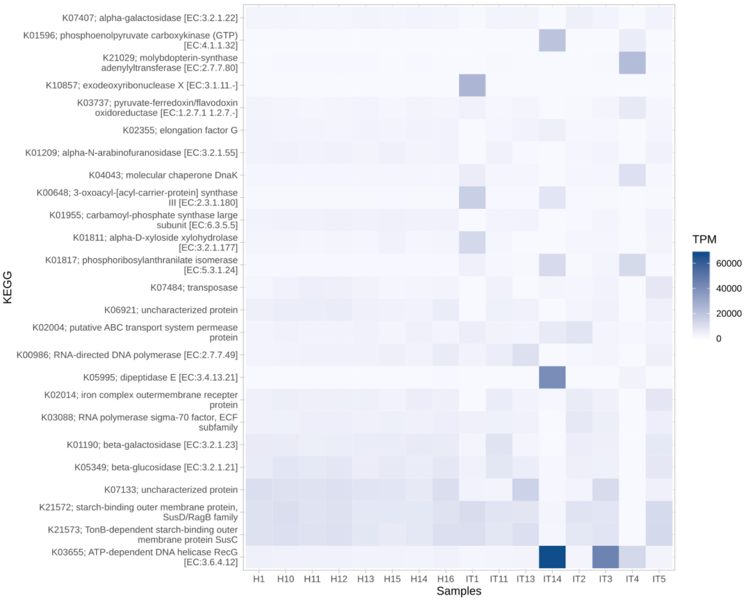

**Figure S12:** Most abundant KEGGs in the 'prevotella' subset. The abundance of each function (KEGG) is counted in TPM, genes from each function per million genes in the subsetted taxon (*Prevotella*).

**Exploring a specific bin.** E.g. 'maxbin.005.fasta.contigs'.

```
maxbin005 = subsetBins(Hadza, 'maxbin.005.fasta.contigs' )
# Plot functions
plotFunctions(maxbin005, fun_level = 'KEGG', count = 'tpm')
```

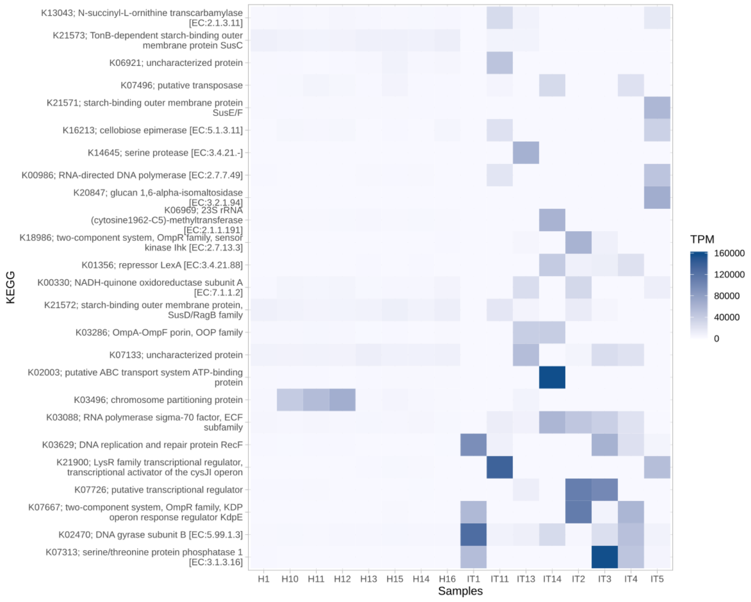

**Figure S13:** Most abundant KEGGs in the 'maxbin005' subset. The abundance of each function (KEGG) is counted in TPM, genes from each function per million genes in the subsetted bin.

**Combining different subsets.** E.g. Carbohydrate metabolism of *Prevotella* and *Bacteroides*

```
# Subset ORFs related to carbohydrate-metabolism
carbohydrates = subsetFun(Hadza, 'Carbohydrate metabolism' )
# From 'Carbohydrate metabolism' subset 'Prevotella'
prevotella_carbohydrates = subsetTax(carbohydrates, 'genus',  'Prevotella')
# From 'Carbohydrate metabolism' subset 'Bacteroides'
bacteroides_carbohydrates = subsetTax(carbohydrates,  'genus','Bacteroides')
# Combine both SQM objects
carbohydrates_prevotella_bacteroides = combineSQM(prevotella_carbohydrates,
               bacteroides_carbohydrates, tax_source =  'contigs')
# Plot taxonomy
plotTaxonomy(carbohydrates_prevotella_bacteroides, 'genus', 'percent', N = 10,
               rescale= F, others= T)
```

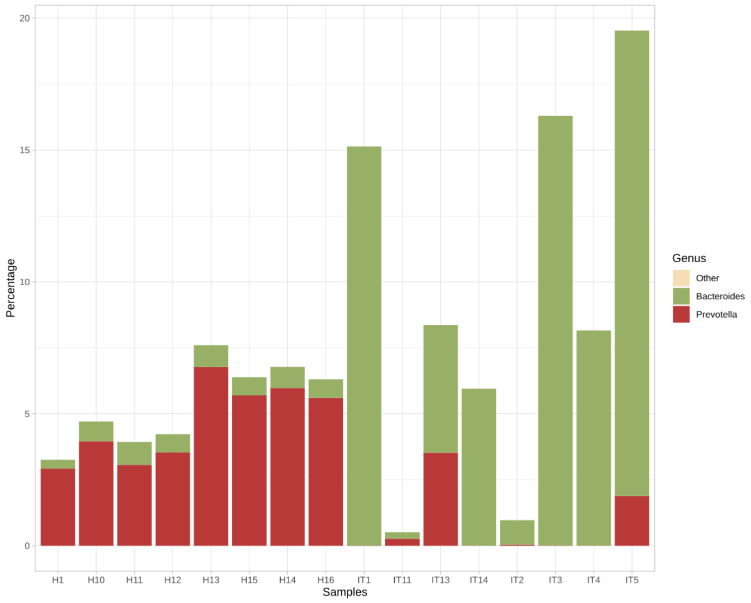

**Figure S14:** Taxonomy barplot at the genus level of the combined object: 'Carbohydrate metabolism of *Bacteroides* and *Prevotella*'. The abundance percentage of each taxonomic group is relative to the whole metagenome.

```
plotFunctions(carbohydrates_prevotella_bacteroides, fun_level = 'KEGG', count = 'tpm')
```

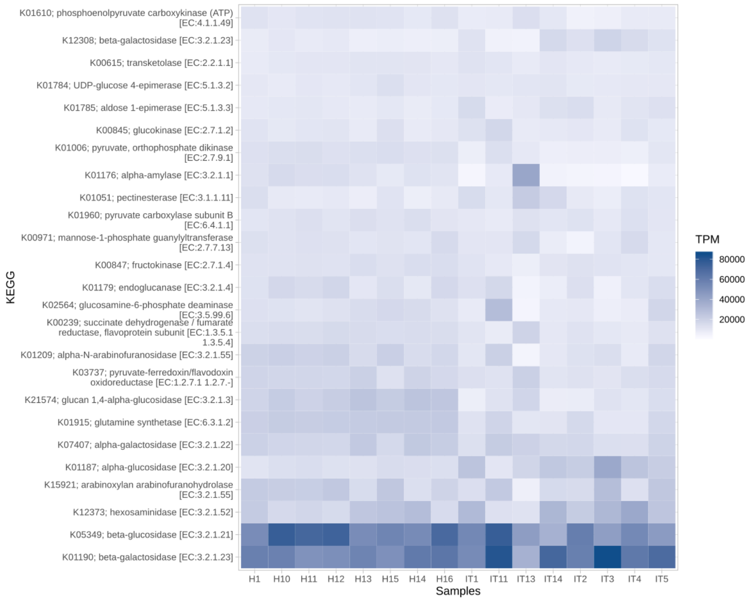

**Figure S15:** Most abundant KEGGs in the combined object: 'Carbohydrate metabolism of *Bacteroides* and *Prevotella*'. Abundances are counted in TPM, genes of each function per million genes in the combine object.

##### 2.2.4. Exporting results: The *exportTable*, *exportKrona* and *exportPathway* functions

###### Export a table

Export results in 'tsv' format, which is supported by software like *Excel* or *LibreOffice Calc*:

```r
exportTable(carbohydrates_prevotella_bacteroides$functions$COG$tpm, 'Hadza_COG_tpm.tsv')
```

###### Export a Krona chart

Also, Krona charts can be exported with ease:

```r
exportKrona(Hadza)
```

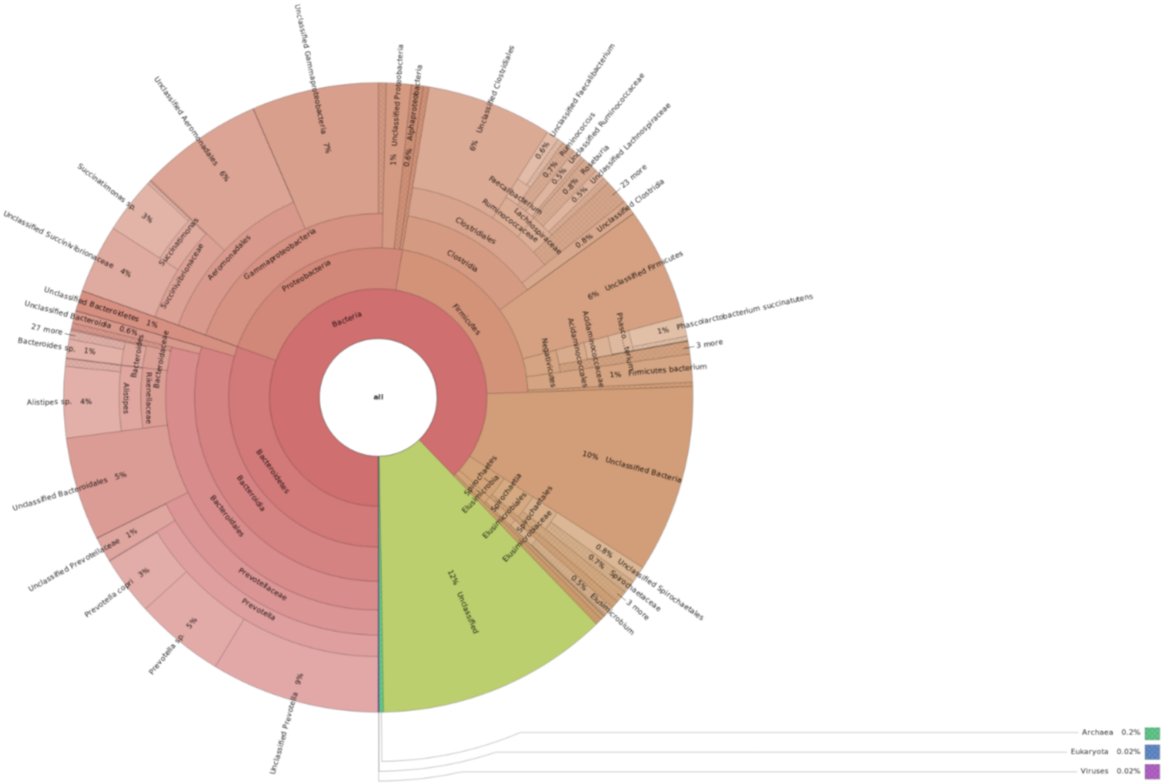

**Figure S16:** Snapshot of Krona chart.

##### Export any KEGG Pathway

Other advantageous feature is the option to analyze whether a KEGG pathway is complete in one or all the samples.

E.g. ko00400 'Phenylalanine, tyrosine and tryptophan biosynthesis':

```
# To know if a KEGG is present or not in one sample,
# the user can provide a vector of colors.
# Colors must be in the same order that samples
# To distinguish between Hadza and Italians:
# Repeat one color 8 times because the first 8 samples are from Hadza.
# The next 8 samples are from Italians, repeat their color.
colors = c(rep('#006682', 8), rep('#c26e00', 8))
exportPathway(Hadza, '00400', output_suffix =  'aromatic_aa', sample_colors =  colors,
              max_scale_value = 3, count ='copy_number')
```

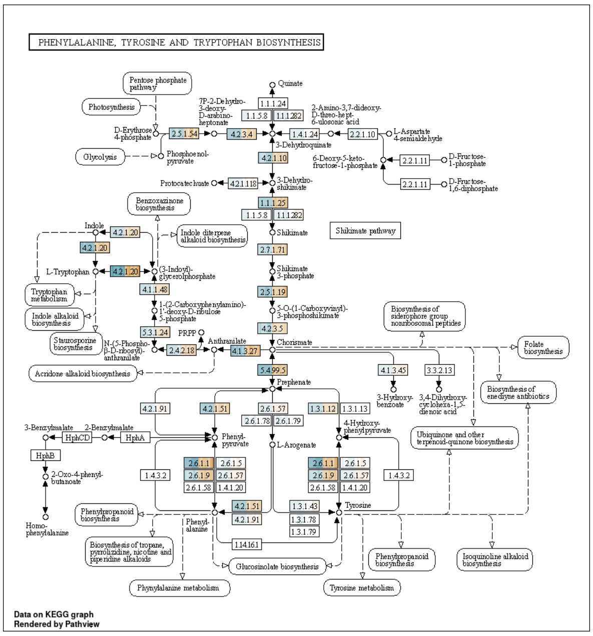

**Figure S17:** Phenylalanine, tyrosine and tryptophan biosynthesis. Each square is an enzyme and it is divided in so many parts as samples are in the metagenome. Each enzyme is linked to one or more KEGGs. If the enzyme is present in one sample, its division is colored with the color of that sample. Hadza samples are shown in blue and Italian ones in orange. The hue of the color depends on the value: the darkest colors imply a copy number $\geq 3$. White is for not present enzymes.

It is also possible to plot the log fold change of TPM or copy number: E.g. ko00400 'Phenylalanine, tyrosine and tryptophan biosynthesis':

```
# To calculate the log-fold change of a KEGG,
# the user can provide a list of vectors.
# One vector with the samples names by condition
#Hadza samples
H.samples = Hadza$misc$samples[grep('H', Hadza$misc$samples)]
IT.samples = Hadza$misc$samples[grep('IT', Hadza$misc$samples)]
condition = list(H.samples, IT.samples)
# Choose one color per condition
colors = c('#006682', '#c26e00')
exportPathway(Hadza, '00400', output_suffix =  'aromatic_aa.log2FC',
     fold_change_colors =  colors, fold_change_groups =condition, count ='copy_number',
     max_scale_value = 1.5)
```

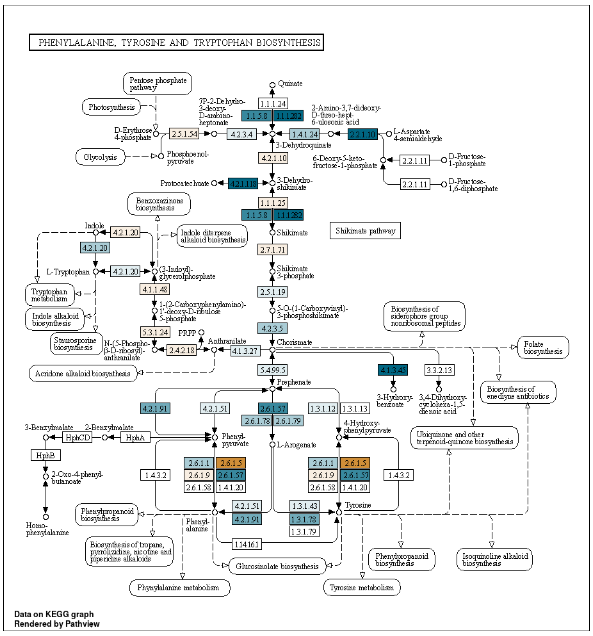

**Figure S18:** Phenylalanine, tyrosine and tryptophan biosynthesis. Each square is an enzyme. The hue of the color depends on the value of the log-fold-change calculated using copy numbers: the darkest orange implies a log-fold-change value ≥ 1.5 (more abundant in Italian samples) and the darkest blue a log-fold-change ≤ -1.5 (more abundant in Hadza samples). White is for not present enzymes.

##### 2.2.5. Using other R packages

Moreover, the user can effortlessly make statistical tests with other R packages using the structure of the SQM Object. For instance:

**Differential abundances with *DESeq2*** E.g. Which KEGGs are significantly more abundant in the Hadza gut microbiome rather than in the Italian one?

```
# Tutorial based on
# http://bioconductor.org/packages/devel/bioc/vignettes/DESeq2/inst/doc/DESeq2.html
# Load the package
library('DESeq2')
# Required: matrix with the raw abundances, colData (metadata file)
# Metadata is a data.frame whose rownames are the sample names, and
# each column is a vector of factors.
# E.g. In this example, the factor (condition) is the country: Hadza or Italy
metadata = as.data.frame(c(rep('Hadza', 8), rep('Italy', 8)))
rownames(metadata) = colnames(Hadza$functions$KEGG$abund)
colnames(metadata) = 'condition'
# Verify sample order
all(rownames(metadata) == colnames(Hadza$functions$KEGG$abund))
# Convert your data to the format required by DESeq2
dds = DESeqDataSetFromMatrix(countData = Hadza$functions$KEGG$abund, colData =
      metadata, design = ~condition)
# Remove low abundant KEGGs:
keep = rowSums(counts(dds)) >= 10
```

```r
dds = dds[keep, ]
# Choose factor levels
dds$condition = factor(dds$condition, levels = c('Hadza', 'Italy'))
# Run DESeq2
dds2 = DESeq(dds)
results = results(dds2)
results05 = results(dds2, alpha = 0.05)
plotMA(results05, ylim = c(-2, 2))
```

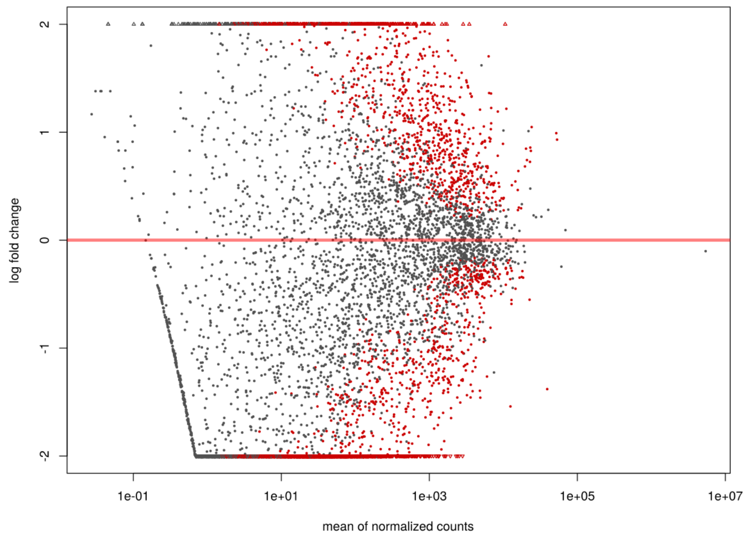

**Figure S19:** Genes differential abundance. Red colored points mean that the adjusted p adjusted value is less than 0.05. Points which fall out of the window are plotted as open triangles pointing either up or down.

Analyzing the 'Starch and sucrose metabolism' in the Hadza and Italian gut microbiome:

```r
# Select significantly most abundant KEGGs in the Hadza samples
H.up = rownames(results05)[!is.na(results05$padj) &
     results05$padj < 0.05 & results05$log2FoldChange < (-1)]
# Negative FC: enriched in the "control".
# log2FC <(-1): at least 2 times more abundant in the "control" .
# Select significantly most abundant KEGGs in the Italian samples
IT.up = rownames(results05)[!is.na(results05$padj)
& results05$padj < 0.05 & results05$log2FoldChange > 1]
# KO terms from a given functional category. E.g. starch and sucrose metabolism
KO.ss = names(Hadza$misc$KEGG_path)[grepl('Starch and sucrose metabolism',
Hadza$misc$KEGG_path, fixed=T)]
# Select KEGGs from the starch and sucrose metabolism KEGG pathway
that have a higher abundance in the Hadza samples
H.up.ss = intersect(H.up, KO.ss)
# Show examples
```

```r
Hadza$misc$KEGG_names[H.up.ss]
[K00694] "cellulose synthase (UDP-forming) [EC:2.4.1.12]"
[K00697] "trehalose 6-phosphate synthase [EC:2.4.1.15 2.4.1.347]"
[K00844] "hexokinase [EC:2.7.1.1]"
[K01087] "trehalose 6-phosphate phosphatase [EC:3.1.3.12]"
[K01194] "alpha,alpha-trehalase [EC:3.2.1.28]"
[K02438] "glycogen debranching enzyme [EC:3.2.1.196]"
[K13057] "trehalose synthase [EC:2.4.1.245]"
[K15778] "phosphomannomutase / phosphoglucomutase [EC:5.4.2.8 5.4.2.2]"
[K15916] "glucose/mannose-6-phosphate isomerase [EC:5.3.1.9 5.3.1.8]"
[K19668] "cellulose 1,4-beta-cellobiosidase [EC:3.2.1.91]"
[K07405] "alpha-amylase [EC:3.2.1.1]"
[K16149] "1,4-alpha-glucan branching enzyme [EC:2.4.1.18]"
# Select KEGGs from the starch and sucrose metabolism KEGG pathway
that have a higher abundance in the Italian samples
IT.up.ss = intersect(IT.up, KO.ss)
# Show examples
Hadza$misc$KEGG_names[IT.up.ss]
[K00689] "dextransucrase [EC:2.4.1.5]"
[K00691] "maltose phosphorylase [EC:2.4.1.8]"
[K01178] "glucoamylase [EC:3.2.1.3]"
[K01193] "beta-fructofuranosidase [EC:3.2.1.26]"
[K01210] "glucan 1,3-beta-glucosidase [EC:3.2.1.58]"
[K07024] "sucrose-6-phosphatase [EC:3.1.3.24]"
[K16055] "trehalose 6-phosphate synthase/phosphatase [EC:2.4.1.15 3.1.3.12]"
```

It is easy to check individually the abundance of some of these KEGGs. For instance, we show a box plot of the abundances of the KEGG K19668 which corresponds to the "cellulose 1,4-beta-cellobiosidase [EC:3.2.1.91]":

```r
# Extract TPM values for K19668 in Hadza and Italians samples
H.K19668 = Hadza$functions$KEGG$tpm['K19668',
               grep('H', colnames(Hadza$functions$KEGG$tpm))]
IT.K19668 = Hadza$functions$KEGG$tpm['K19668',
               grep('IT', colnames(Hadza$functions$KEGG$tpm))]
# Boxplot
boxplot(H.K19668, IT.K19668, names = c('Hadza', 'Italy'), main = 'K19668',
        ylab = 'TPM', outline = F, col = c('#006682', '#c26e00'))
```

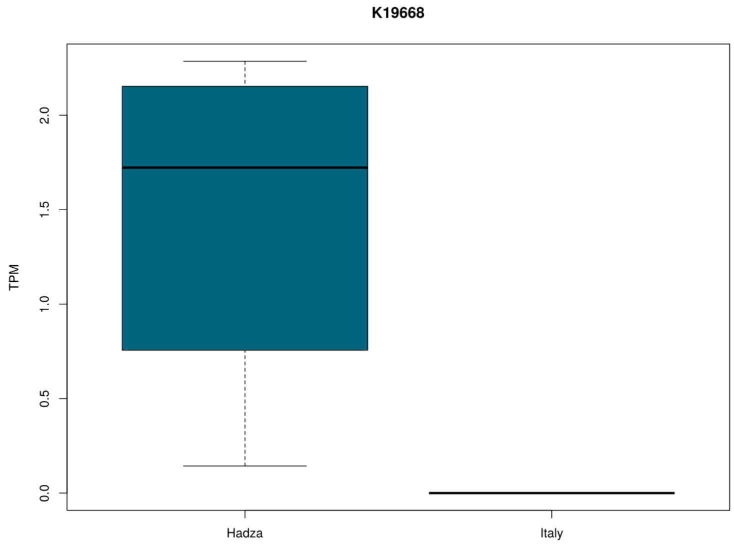

**Figure S20:** Boxplot of the abundance (TPM) of the KEGG K19668 in the Hadza and Italian samples.

E.g. Which taxa at the family rank are significantly more abundant in the Hadza gut microbiome rather than in the Italian one?

```
# Like with KEGGs, load the package
library('DESeq2')
# Required: matrix with the raw abundances, colData (metadata file)
metadata = as.data.frame(c(rep('Hadza', 8), rep('Italy', 8)))
rownames(metadata) = colnames(Hadza$taxa$family$abund)
colnames(metadata) = 'condition'
# Verify sample order
Now:
all(rownames(metadata) == colnames(Hadza$taxa$family$abund))
# Convert your data to the format required by DESeq2
dds = DESeqDataSetFromMatrix(countData = Hadza$taxa$family$abund, colData =
metadata, design = ~condition)
# Remove low abundant taxa:
keep = rowSums(counts(dds)) >= 10
dds = dds[keep, ]
# Choose factor levels
dds$condition = factor(dds$condition, levels = c('Hadza', 'Italy'))
# Run DESeq2
dds2 = DESeq(dds)
results = results(dds2)
results05 = results(dds2, alpha = 0.05)
plotMA(results05)
```

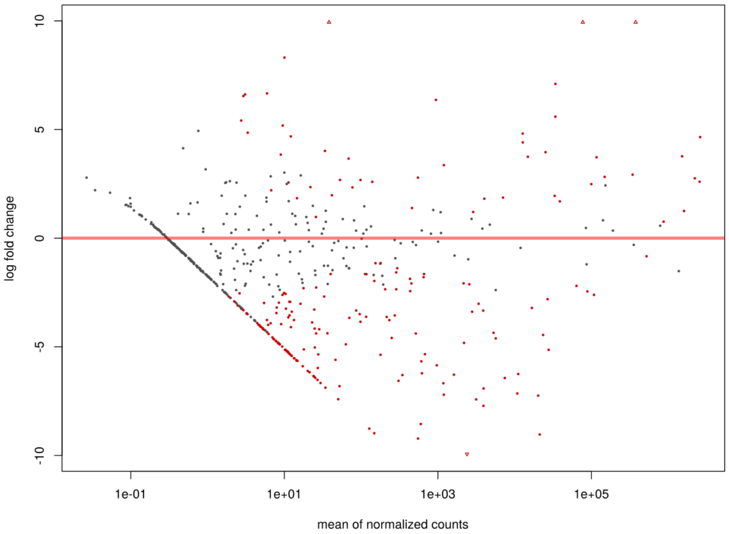

**Figure S21:** Taxa differential abundance. Red colored points mean that the adjusted p adjusted value is less than 0.05. Points which fall out of the window are plotted as open triangles pointing either up or down.

```
# Select taxa that have a higher abundance in the Hadza samples
H.up = rownames(results05)[!is.na(results05$padj) & results05$padj < 0.05
& results05$log2FoldChange < (-1)]
# Show results
H.up
[1] "Unclassified Candidatus Bathyarchaeota"
[2] "Unclassified Candidatus Lokiarchaeota"
[3] "Unclassified Candidatus Thorarchaeota"
[4] "Unclassified Methanobacteriales"
[5] "Unclassified Methanomicrobiales"
[6] "methanogenic archaeon (no family in NCBI)"
[7] "Methanosarcinaceae"
[8] "Unclassified Methanomicrobia"
[9] "Candidatus Methanomethylophilaceae"
[10] "Methanomassiliicoccaceae"
# Select taxa that have a higher abundance in the Italian samples
IT.up = rownames(results05)[!is.na(results05$padj) & results05$padj < 0.05 &
 results05$log2FoldChange > 1]
# Show examples
IT.up
[1] "Actinomycetaceae"  "Bifidobacteriaceae"
[3] "Unclassified Bifidobacteriales"  "Corynebacteriaceae"
[5] "Nocardiaceae"  "Propionibacteriaceae"
[7] "Coriobacteriaceae" "Eggerthellaceae"
[9] "Unclassified Coriobacteriia" "Bacteroidaceae"
```

##### Using the *vegan* package

How similar are the samples among them if we look at the genus taxonomic rank? We run a NMDS (Non metric dimensional scaling) to answer this question.

```
library('vegan')
# Metadata is a data.frame whose rownames are the sample names, and
# each column is a vector of factors.
# E.g. In this example, the factor (condition) is the country: Hadza or Italy
metadata = as.data.frame(c(rep('Hadza', 8), rep('Italy', 8)))
rownames(metadata) = colnames(Hadza$taxa$genus$percent)
colnames(metadata) = 'condition'
# Tranpose the matrix to have samples in rows.
genus_pc = t(Hadza$taxa$genus$percent)
MDS = metaMDS(genus_pc)
colvec = c('#006682','#c26e00') # colors
plot(MDS, display = 'sites')
with(metadata, points(MDS, display = 'sites', col = colvec[condition], pch = 19))
with(metadata, legend('topright', legend = levels(condition), col = colvec, pch = 21,
pt.bg = colvec))
```

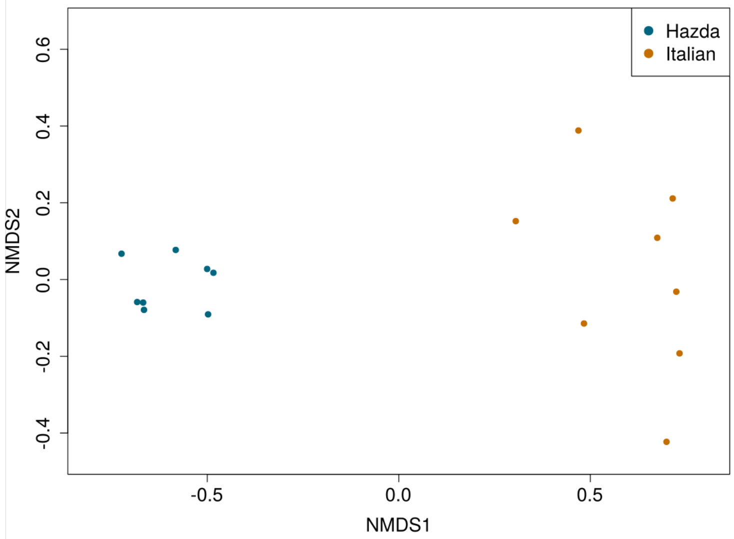

**Figure S22:** Non-metric multidimensional scaling (NMDS) ordination at the genus rank.

Additionally, we ran a PERMANOVA (Permutational Multivariate Analysis Of Variance Using Distance Matrices) test to know if the country affects the clustering of the samples in the previous ordination:

```
country = c(rep('Hadza', 8), rep('Italy', 8))
# Run adonis to make a PERMANOVA in the vegan package
adonis(genus_pc~country)
Call :
adonis (formula = genus_pc ~ country)
```

```
Permutation : free
Number of permutation: 999
Terms added sequentially (first to last)
          Df SumsOfSqs MeanSqs F.Model      R2 Pr(>F)
country    1   0.85836 0.85836  17.107 0.54995  0.001 ***
Residuals 14   0.70245 0.05017         0.45005
Total     15   1.56081                 1.00000
---
Signif. codes:  0 '***' 0.001 '**' 0.01 '*' 0.05 '.' 0.1 ' ' 1
```

From this result, we can conclude that the samples from the same country are more similar among them, when we analyze the taxonomical distribution at the genus rank.

How similar are the samples among them if we look the functional annotations?

```
library('vegan')
metadata = as.data.frame(c(rep('Hadza', 8), rep('Italy', 8)))
rownames(metadata) = colnames(Hadza$functions$KEGG$tpm)
colnames(metadata) = 'condition'
# Tranpose the matrix to have samples in rows.
kegg_tpm = t(Hadza$functions$KEGG$tpm)
MDS = metaMDS(kegg_tpm)
colvec = c('#006682','#c26e00') # colors
plot(MDS, display = 'sites')
with(metadata, points(MDS, display = 'sites', col = colvec[condition], pch = 19))
with(metadata, legend('topright', legend = levels(condition), col = colvec, pch = 21,
pt.bg = colvec))
```

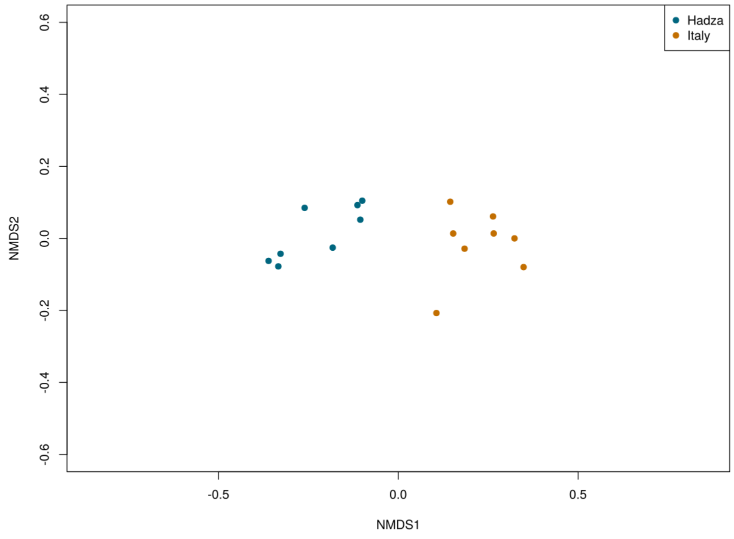

**Figure S23:** Non-metric multidimensional scaling (NMDS) ordination using KEGGs.

We ran a PERMANOVA test to know if the country is affecting the clustering of samples in the ordination:

```
country = c(rep('Hadza', 8), rep('Italy', 8))
adonis(kegg_tpm~country)
Call :
adonis (formula = kegg_tpm ~ country)
Permutation : free
Number of permutation: 999
Terms added sequentially (first to last)
          Df  SumsOfSqs  MeanSqs F.Model      R2 Pr(>F)
country    1  0.034942 0.034942  7.9894 0.36333  0.001 ***
Residuals 14  0.061231 0.004374         0.63667
Total     15  0.096173                  1.00000
---
Signif. codes:  0 '***' 0.001 '**' 0.01 '*' 0.05 '.' 0.1 ' ' 1
```

According to this, samples from the same country are more similar in their functional annotations (KEGGs).

#### 3. Visualising a SqueezeMeta project in the anvi'o platform

The second option to explore a SqueezeMeta project is the use of the anvi'o platform (http://merenlab.org/software/anvio/):

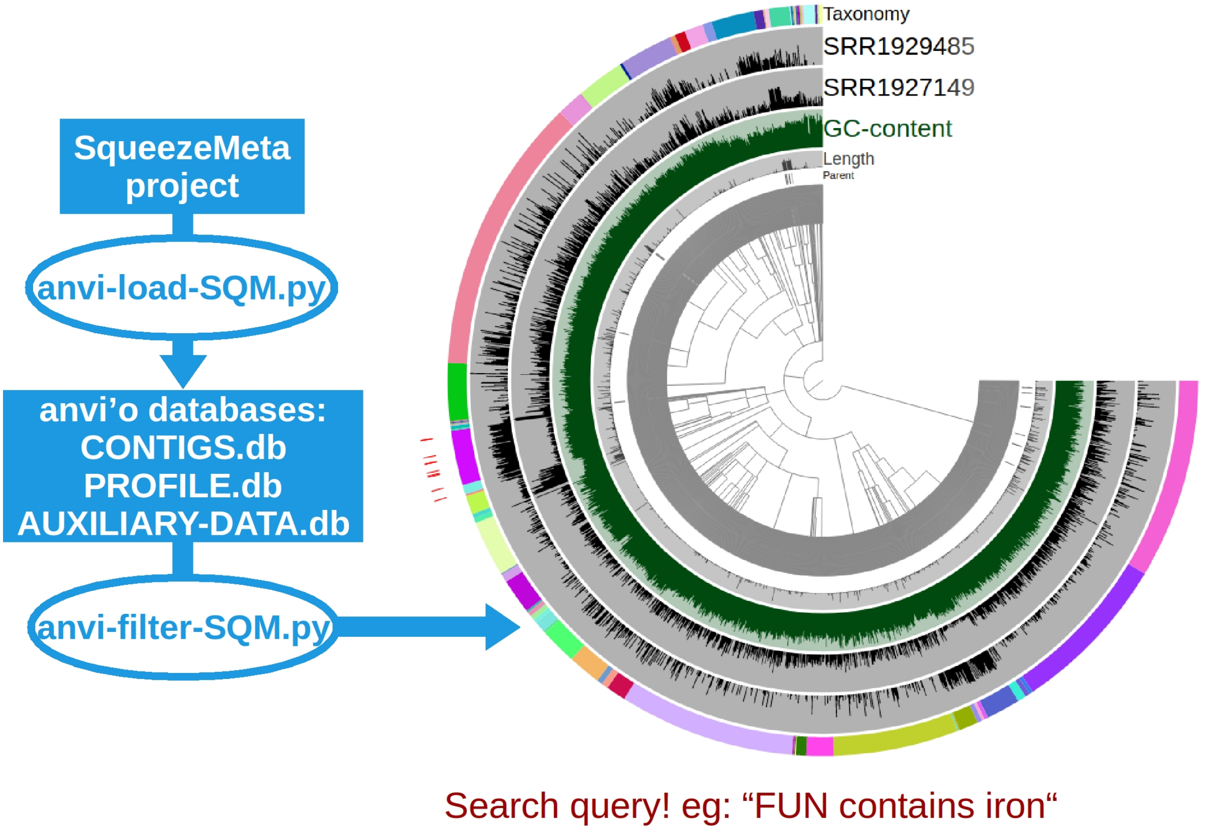

**Figure S24:** Anvi'o workflow using SQMtools.

##### 3.1. Activating the anvi'o environment

The activation of the anvi'o environment will depend on the installation procedure. Follow their instructions: http://merenlab.org/2016/06/26/installation-v2/ to be sure it is working properly.

```
conda activate anvio-6.1# anvio-6.1 version
```

##### 3.2. Parsing a SqueezeMeta project into anvi'o

In order to load a SqueezeMeta project into anvi'o it is necessary to run the *anvi-load-sqm.py* script:

```
python3 /path/to/SqueezeMeta/utils/anvio_utils/anvi-load-sqm.py -p Hadza16
-o Hadza16/results/anvio --num-threads 12 --min-contig-length 2500
--run-scg-taxonomy --run-HMMS
```

where:

- **p** : SqueezeMeta project directory
- **o** : Output directory where anvi'o files will be created
- **run-HMMS** : Run anvi-run-hmms command of anvi'o for identifying single-copy core genes
- **run-scg-taxonomy** : Run anvi-run-scg-taxonomy command of anvi'o for affiliating single-copy core genes with taxonomic names
- **min-contig-length** : Minimum length of the contigs that will be profiled by anvi'o.
- **num-threads** : Number of threads

At this point, the whole SqueezeMeta project will be formatted as anvi'o requires. It saves the information of contigs, genes, and bins into databases (CONTIGS.db and PROFILE.db). One of the most beneficial advantages of using anvi'o is that it gathers all this information in an interactive interface. With this interface, the user is able to immediately visualise the 'splits' (fragments of contigs that anvi'o splits because they are larger than a certain size), their coverage, their taxonomy and their genes and functions.

#### 3.3. Filtering data from a SqueezeMeta project

Here, we show two different examples on how to analyze a SqueezeMeta project with anvi'o using the *anvi-filter-sqm.py* script (description of how to make the queries is in the documentation of the script).
**Example 1:**
Picking up all the contigs that have genes whose annotations include the word 'antibiotic':

```
/path/to/SqueezeMeta/utils/anvio_utils/anvi-filter-sqm.py
-c Hadza16/results/anvio/CONTIGS.db -p Hadza16/results/anvio/PROFILE.db
-t Hadza16/results/anvio/Hadza16_anvio_contig_taxonomy.txt
-o Hadza16/results/anvio/antibiotic -q '(FUN CONTAINS antibiotic)'
```

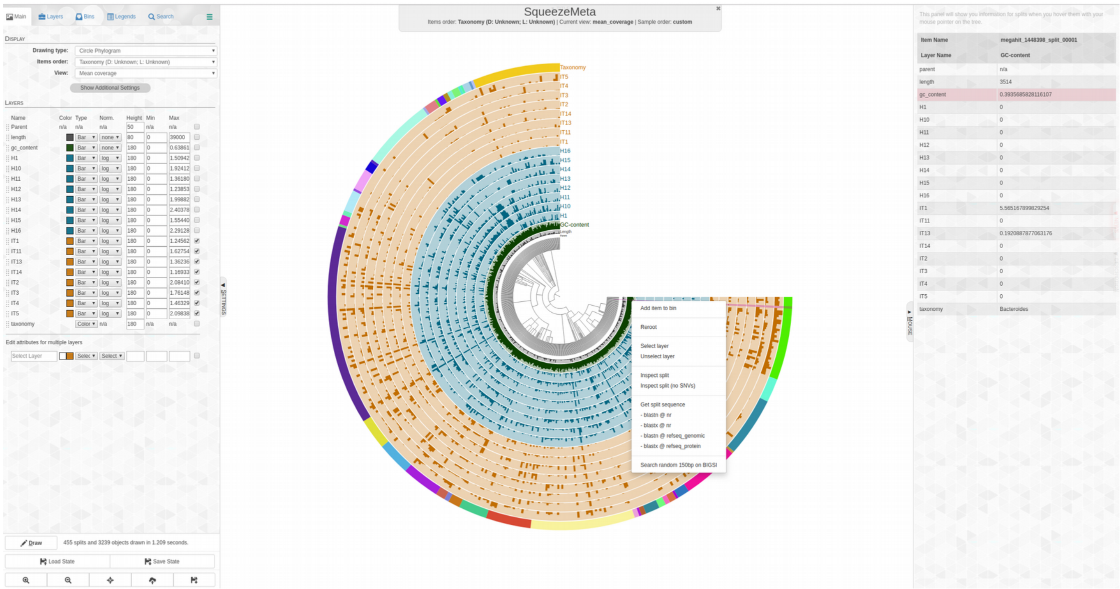

**Figure S25:** View of the anvi-interactive mode. The graph shows a general view of all the splits in the project. The right panel allows the user to control some features of the interface such as the splits' order, the colors and the layer sizes. The left panel indicates, per split, its abundance in the different samples and its taxonomy. Look http://merenlab.org/2016/02/27/the-anvio-interactive-interface/ for more information.

**Example 2:**

Exploring splits related with the '00400 Phenylalanine, tyrosine and tryptophan biosynthesis' KEGG Pathway :

```
/path/to/SqueezeMeta/utils/anvio_utils/anvi-filter-sqm.py
-c Hadza16/results/anvio/CONTIGS.db -p Hadza16/results/anvio/PROFILE.db
-t Hadza16/results/anvio/Hadza16_anvio_contig_taxonomy.txt
-q'(FUNH CONTAINS Phenylalanine, tyrosine and tryptophan biosynthesis)'
-o Hadza16/results/anvio/aromatic_aa_synthesis
```

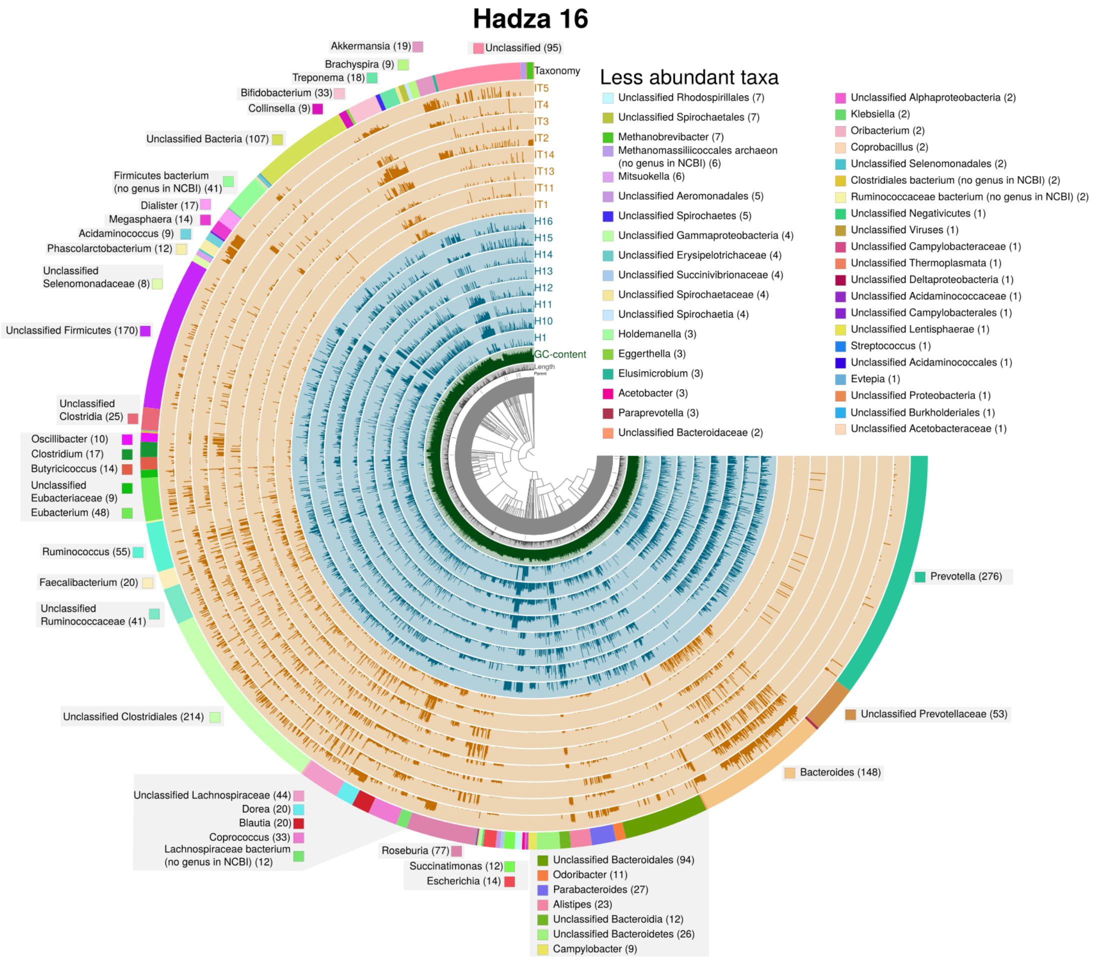

**Figure S26:** View of the splits that contain at least one gene from the '00400 Phenylalanine, tyrosine and tryptophan biosynthesis' KEGG Pathway.

To improve the visualisation, it is possible to make more complex queries. Using the previous query, we also remove all the contigs that have not been classified at the order level:

```
/path/to/SqueezeMeta/utils/anvio_utils/Figureanvi-filter-sqm.py
-c Hadza16/results/anvio/CONTIGS.db -p Hadza16/results/anvio/PROFILE.db
-t Hadza16/results/anvio/Hadza16_anvio_contig_taxonomy.txt -q
'(FUNH CONTAINS 'Phenylalanine, tyrosine and tryptophan biosynthesis'
AND GENUS NOT IN ['Unclassified','Unclassified Firmicutes', 'Unclassified Bacteria',
'Unclassified Clostridiales','Unclassified Spirochaetales', 'Unclassified Clostridia',
'Unclassified Aeromonadales', 'Unclassified Spirochaetes', 'Unclassified Bacteroidia',
'Unclassified Alphaproteobacteria', 'Unclassified Selenomonadales',
'Unclassified Viruses', 'Unclassified Thermoplasmata', 'Unclassified Lentisphaerae',
'Unclassified Deltaproteobacteria','Unclassified Campylobacterales',
'Unclassified Acidaminococcales','Unclassified Proteobacteria',
'Unclassified Spirochaetia', 'Unclassified Negativicutes', 'Unclassified Bacteroidales',
```

```
'Unclassified Bacteroidetes', 'Unclassified Rhodospirillales',
'Unclassified Burkholderiales', 'Unclassified Gammaproteobacteria'])'
-o Hadza16/results/anvio/aromatic_aa_synthesis_noUnclassified
```

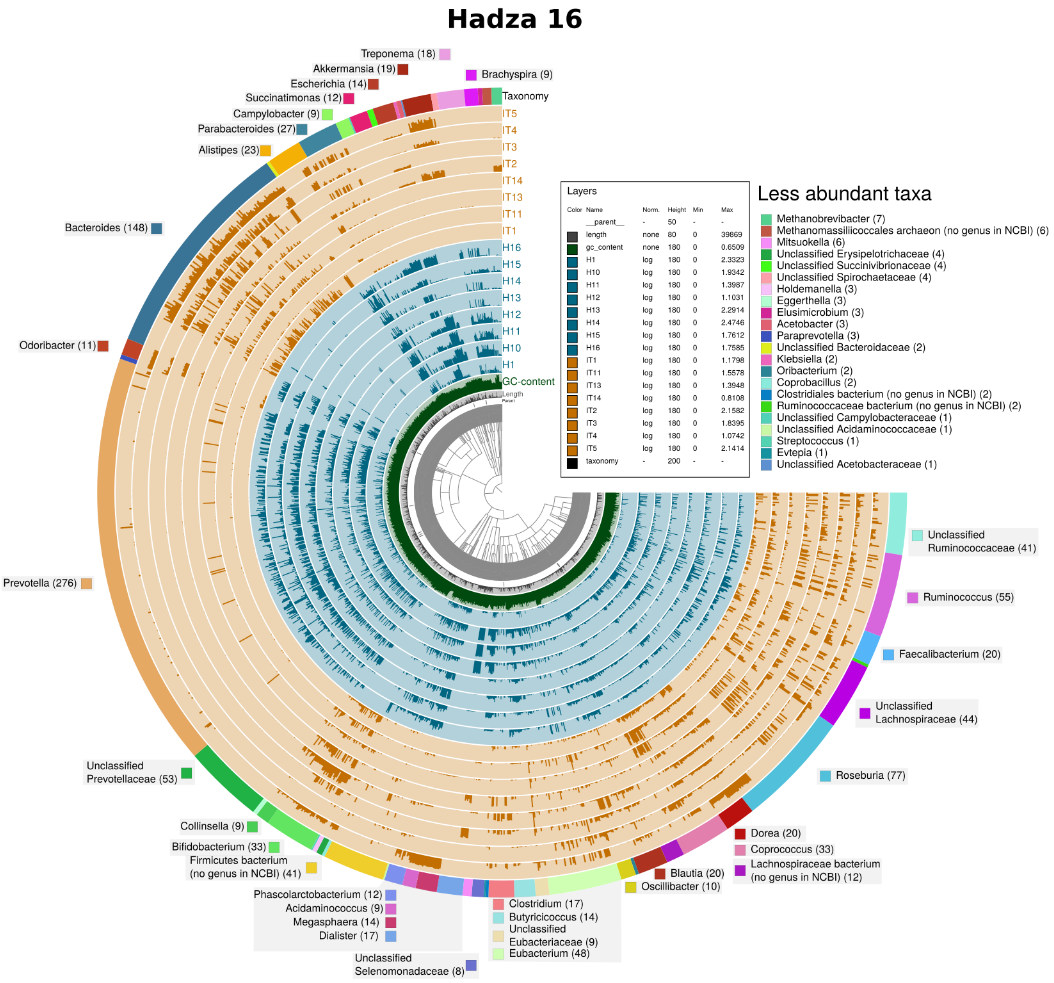

**Figure S27:** View of the splits that contain at least one gene from the '00400 Phenylalanine, tyrosine and tryptophan biosynthesis' KEGG Pathway and have a taxonomic annotation at the order level.

Using the interactive interface it is possible to search for functions, taxonomy, abundance within other options:

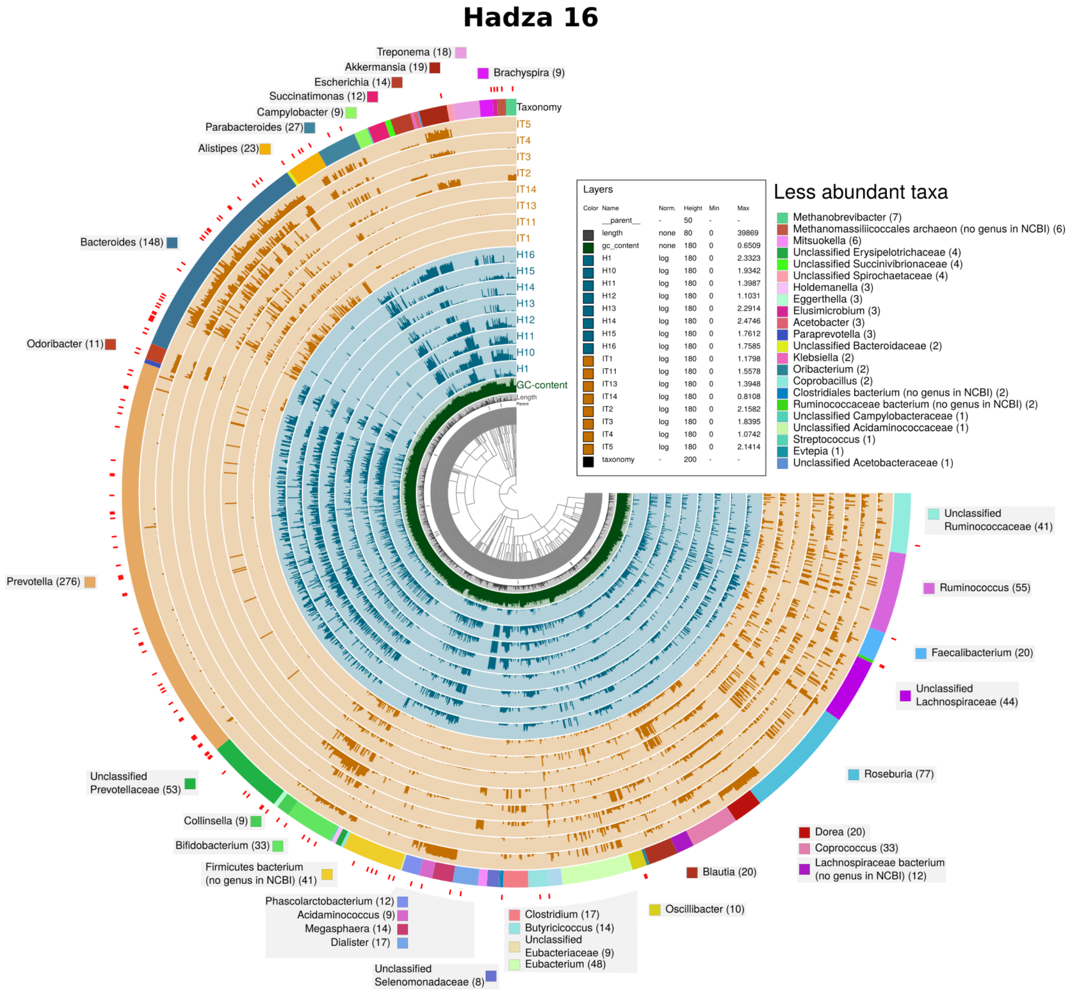

**Figure S28:** The splits highlighted in red contain at least one gene annotated with the KEGG 'K00812'.

#### 3.4. Exploring bins using anvi'o

Anvi'o is a really helpful tool to explore bins. Although, we recommend visiting its tutorials about binning, we decided to show here some of its most handy commands.

The *anvi-interactive* command with the flag *-C <Bin collection name>* is useful to visualise at first glance all the bins of a SqueezeMeta project (it contains one default collection called 'DAS'). This view shows the number of bins, their completitude and redundancy (http://merenlab.org/2016/06/09/assessing-completion-and-contamination-of-MAGs/), and their abundance in each sample:

```
anvi-interactive -c Hadza16/results/anvio/CONTIGS.db
-p Hadza16/results/anvio/PROFILE.db -C DAS
```

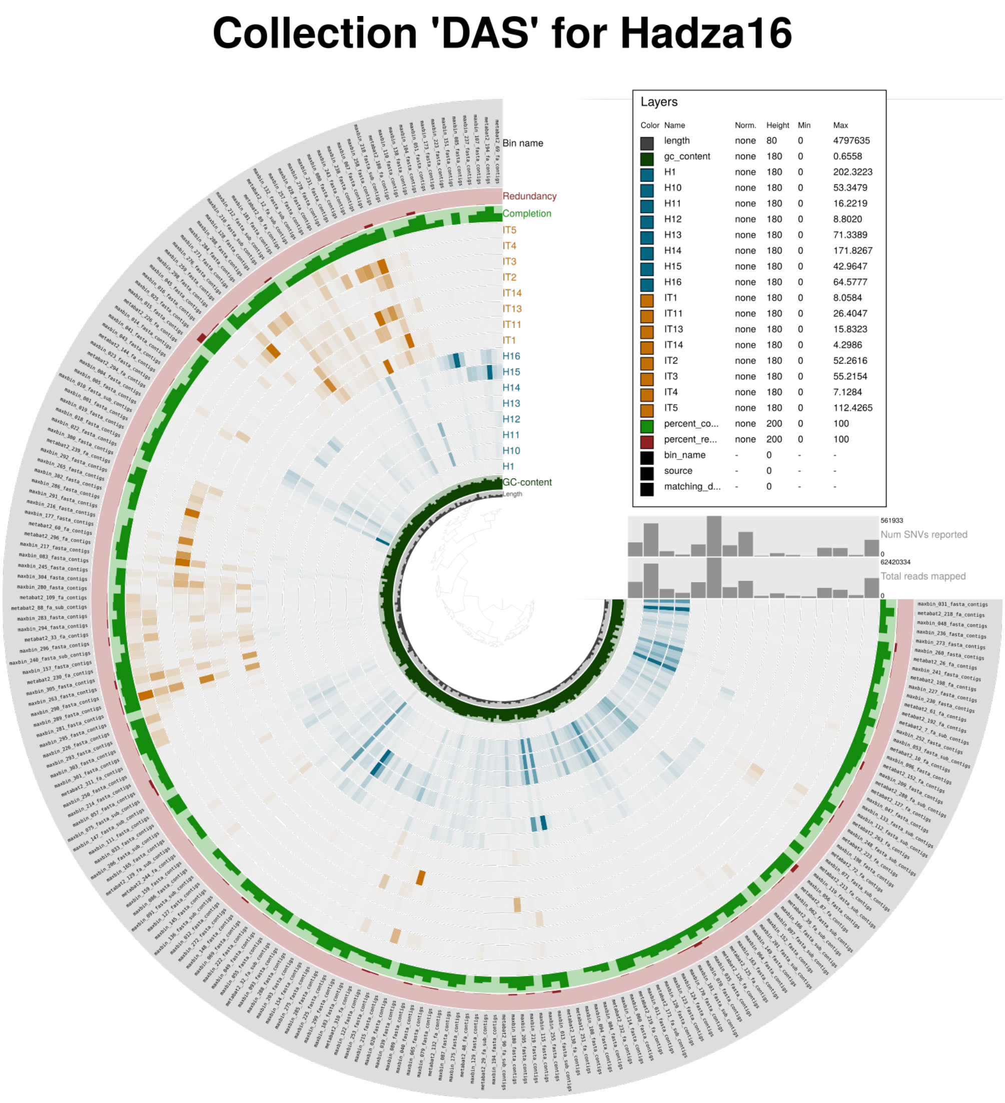

**Figure S29:** General view of all the bins of a SqueezeMeta project.

Moreover, anvi'o allows the user to focus on a specific bin:

```
anvi-refine -c Hadza16/results/anvio/CONTIGS.db -p
Hadza16/results/anvio/PROFILE.db -C DAS -b maxbin_005_fasta_contigs
```

### Refining maxbin_005_fasta_contigs from "DAS"

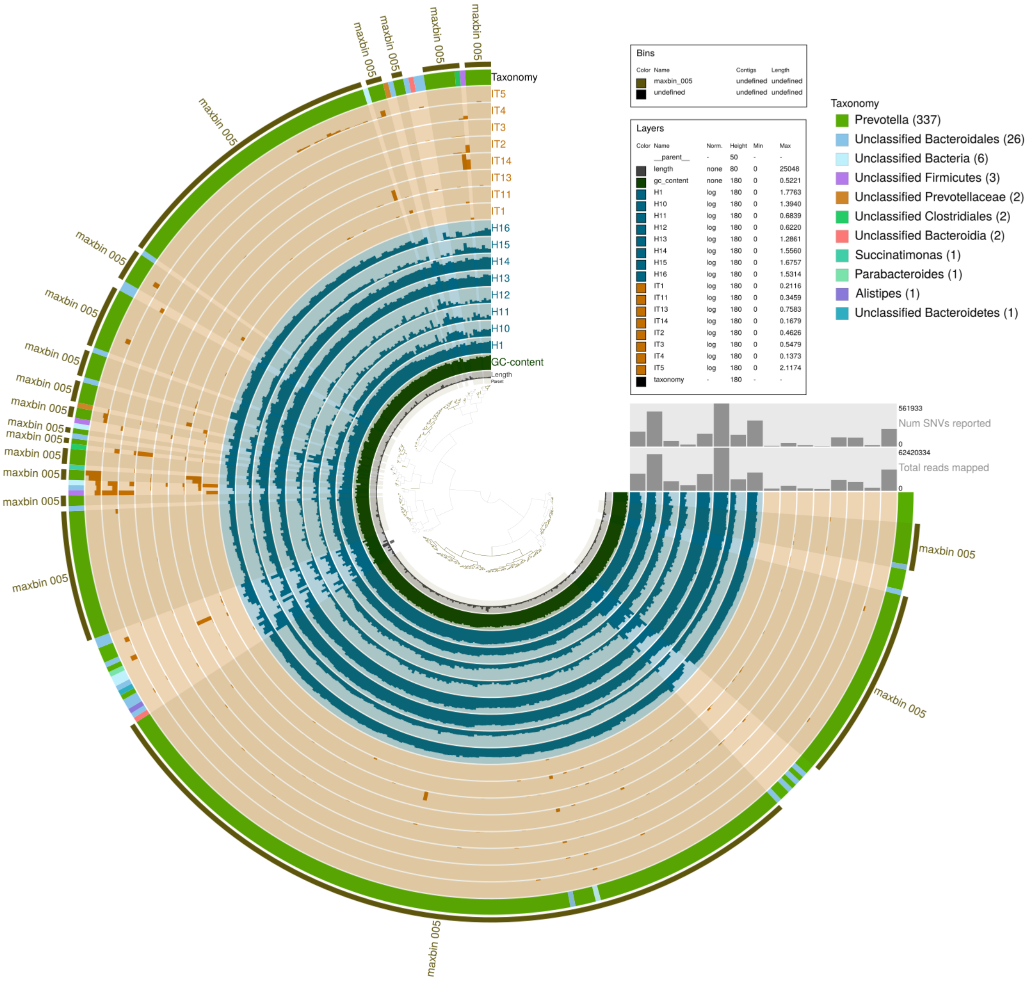

**Figure S30:** Refining a specific bin (‘maxbin_005_fasta_contigs’).
